# LIANA+: an all-in-one cell-cell communication framework

**DOI:** 10.1101/2023.08.19.553863

**Authors:** Daniel Dimitrov, Philipp Sven Lars Schäfer, Elias Farr, Pablo Rodriguez Mier, Sebastian Lobentanzer, Aurelien Dugourd, Jovan Tanevski, Ricardo Omar Ramirez Flores, Julio Saez-Rodriguez

**Affiliations:** Heidelberg University, Faculty of Medicine, and Heidelberg University Hospital, Institute for Computational Biomedicine, Heidelberg, Germany

**Keywords:** Cell-cell communication, Python, framework, spatial, single-cell, multimodal, transcriptomics

## Abstract

The growing availability of single-cell and spatially-resolved transcriptomics has led to the rapidly growing popularity of methods to infer cell-cell communication. Many approaches have emerged, each capturing only a partial view of the complex landscape of cell-cell communication.

Here, we present LIANA+, a scalable framework to decode coordinated inter- and intracellular signalling events from single- and multi-condition datasets in both single-cell and spatially-resolved data. Beyond integrating and extending established methodologies and a rich knowledge base, LIANA+ enables novel analyses using diverse molecular mediators, including those measured in multi-omics data. Accessible as an open-source Python package at https://github.com/saezlab/liana-py, LIANA+ provides a comprehensive set of synergistic components to study cell-cell communication.

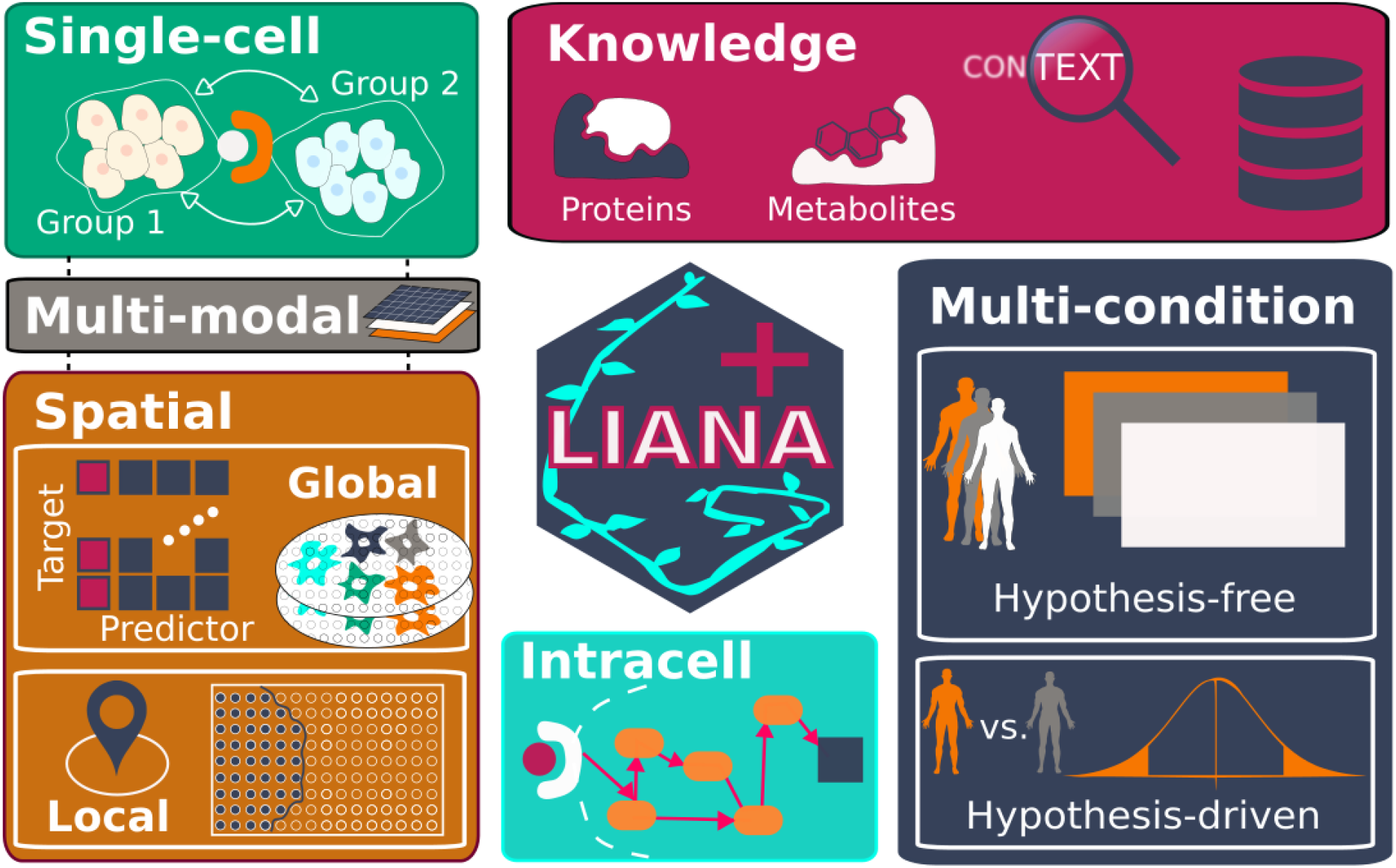

## 1. Background

Cell-cell communication (CCC) inference has recently emerged as a major component of the analysis of single-cell and spatially-resolved transcriptomics data ^1^. Many computational tools have been developed for this purpose, each contributing valuable ideas and developments.

The simplest class of CCC inference methods are those that infer protein-protein interactions from dissociated single-cell transcriptomics data, commonly referred to as ligand-receptor interaction inference methods ^2–5^. Moreover, there are tools that combine intercellular communication with intracellular signalling ^6–10^. All methods are based on multiple assumptions, including the assumption that gene co-expression between dissociated cells, or groups of cells, reflects CCC within tissues ^11^.

In contrast to dissociated single-cell data, spatially-resolved omics technologies preserve spatial context and are thus thought to better reflect the intercellular events that occur within tissues. As a consequence, multiple methods that utilise spatial information have been developed to study CCC ^1^. Typically, these methods infer relationships between proteins ^12,13^ or cell types (cellular neighbourhoods) ^14,15^. Spatially-informed methods differ in the scale at which interactions are inferred, as some infer relationships globally, summarising them across slides as a whole ^12,13,16^, while others do so locally at the individual cell or spot locations within a slide ^17–20^.

The majority of both single-cell and spatially-informed CCC methods have focused on protein-mediated interactions, predominantly from transcriptomics data ^11,21^, and only few methods infer CCC from multi-omics data ^22^. Yet, emerging multi-omics technologies ^23^ are anticipated to provide a more informed view of CCC events, prompting the development of new tools. Furthermore, as a consequence of the almost exclusive use of transcriptomics data, other modes of intercellular signalling, such as small molecule signalling, have been typically ignored ^11^. Recent methods have attempted to infer metabolite-mediated CCC events, again from transcriptomics data ^24–27^, but such inference remains largely limited by the challenges of inferring metabolite abundance from gene expression ^28^.

While early methods analysed CCC in individual samples or single-condition atlases, increasing sample numbers and experimental design complexity have prompted various strategies to extract differential CCC insights. These strategies include methods that (1) require a list of perturbed variables ^29^, (2) consider each variable independently ^17,30^, (3) make use of dimensionality reduction to perform pairwise comparisons between conditions ^4,31^, or (4) jointly model all variables, samples and cell types simultaneously ^32^. Approach (4) can be thought of as modelling coordinated CCC events, and we refer to it as modelling ‘intercellular programmes’ from here on out.

Finally, CCC methods typically rely on pre-existing knowledge ^11,21^. Thus, extensive effort has been put into curating and extending prior knowledge in the context of CCC, with a focus on gathering protein- ^3,4,33^ and, to a lesser extent, metabolite-mediated interactions ^24,26,27,34^. In addition, in some resources, the interactions are further associated with pathways ^4^ or transcriptional regulators ^9,35^, leading to multiple discordant databases and potential inconsistencies caused solely by the choice of resource ^11^.

All these developments have been led by different groups, using various syntaxes and are typically designed for a specific purpose. Here, we introduce LIANA+ as a comprehensive framework that unifies and expands CCC methods and prior knowledge (**Fig. 1**; **Supp. Table 1**). LIANA+ provides eight methods for the inference of CCC from single-cell transcriptomics (**Fig. 1A**) and eight methods for the inference of global and local relationships in spatially-resolved data (**Fig. 1B**), all of which are applicable to multi-omics data. These are further supplemented with four distinct strategies to extract deregulated CCC events in both hypothesis-free and hypothesis-driven manner (**Fig. 1C**). Moreover, CCC events can be connected to intracellular signalling events (**Fig. 1D**) via the use of a rich knowledge base (**Fig. 1E**).

**Figure 1.**
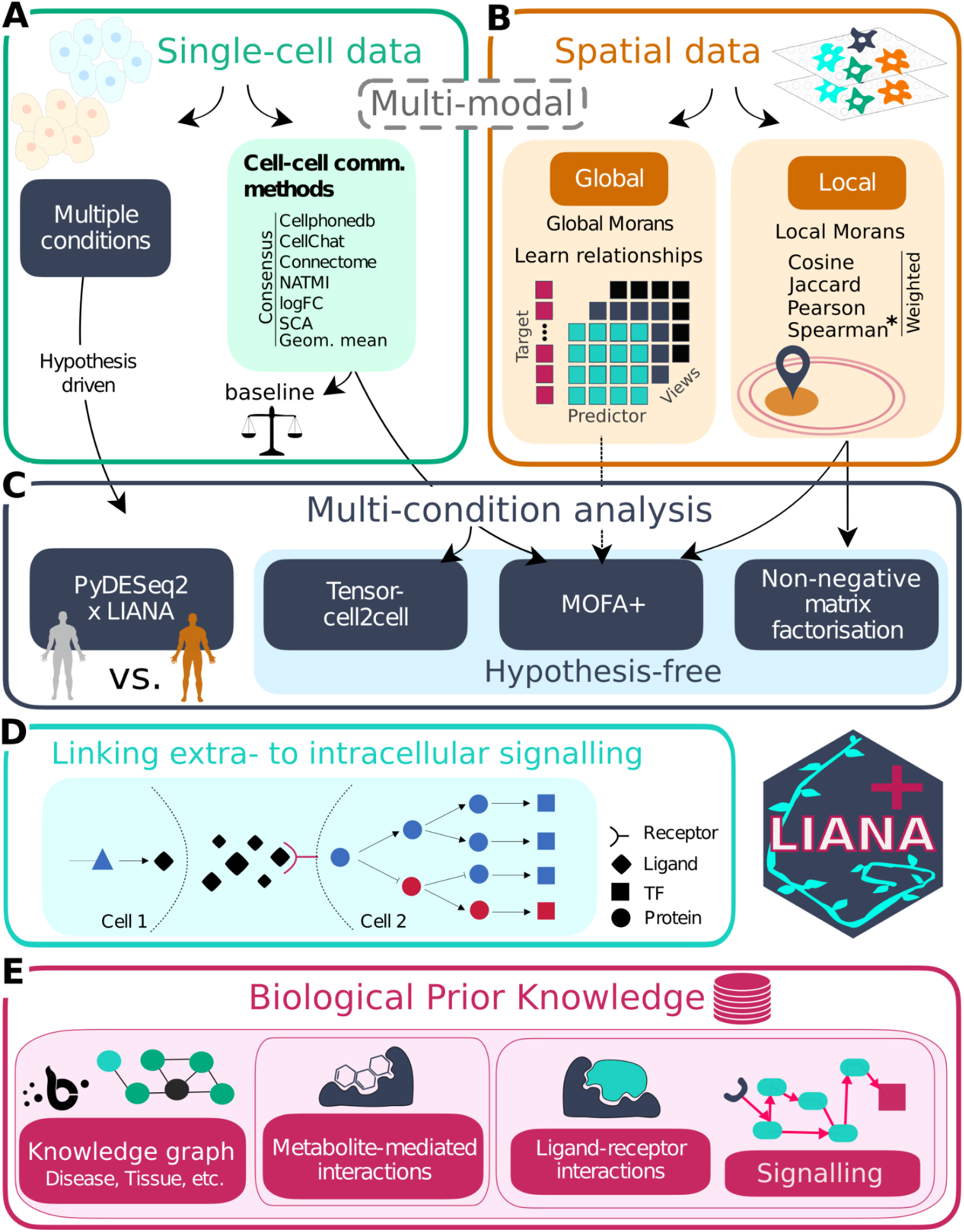
LIANA+ components. Inference of interactions from A) single-cell and B) spatially-resolved (multimodal) omics data. C) CCC analyses across conditions. D) Linking CCC to downstream signalling. E) Extensive prior knowledge.

To showcase the scope and flexibility of LIANA+, we used it to: (1) identify intercellular programmes driving kidney injury response in mouse single-cell and spatial transcriptomics data; (2) extract spatial intercellular patterns in human myocardial infarction; (3) infer CCC using single-cell CITE-seq data from human blood; (4) learn global CCC events and identify the corresponding subregions in spatially-resolved CITE-seq data from human tonsil; (5) identify spatially-informed, metabolite-mediated CCC in the adult mouse brain; (6) find ligand-receptor interactions deregulated in lupus patients; (7) hypothesise intracellular signalling events downstream of CCCs. LIANA+ is an extendable, scverse-compliant ^36^, open-source framework built of synergistic modules, available at https://github.com/saezlab/liana-py, readily applicable to a wide range of single-cell, spatial, multi-omics datasets, with any experimental design.

## 2. Main

### 2.1 LIANA+: an all-in-one framework for cell-cell communication

LIANA+ harmonises current CCC methods and prior knowledge as integral components of the same framework (Fig. 1). All components in LIANA+ use standardised input and output formats, making use of the scverse ecosystem infrastructure ^36^ (**Supp. Fig. S1**). This enables interoperability with external packages and facilitates straightforward and robust extensions of contemporary CCC applications.

#### 2.1.1 LIANA+’s Single-cell Component

In the first iteration of our LIgAnd-receptor aNAlysis (LIANA) framework in R ^11^, we focused on benchmarking existing tools that predict ligand-receptor interactions from dissociated single-cell data. Building on our previous work, LIANA+ is a Python package that natively re-implements eight ligand-receptor methods for the scalable inference of interactions from single-cell data sets (Fig. 2A). These include the algorithms of CellPhoneDB ^3^, CellChat ^4^, Connectome ^5^, NATMI ^37^, SingleCellSignalR ^38^, along with logFC and a geometric mean, as well as their consensus (Fig. 1A**; Supp. Table 2**).

**Figure 2.**
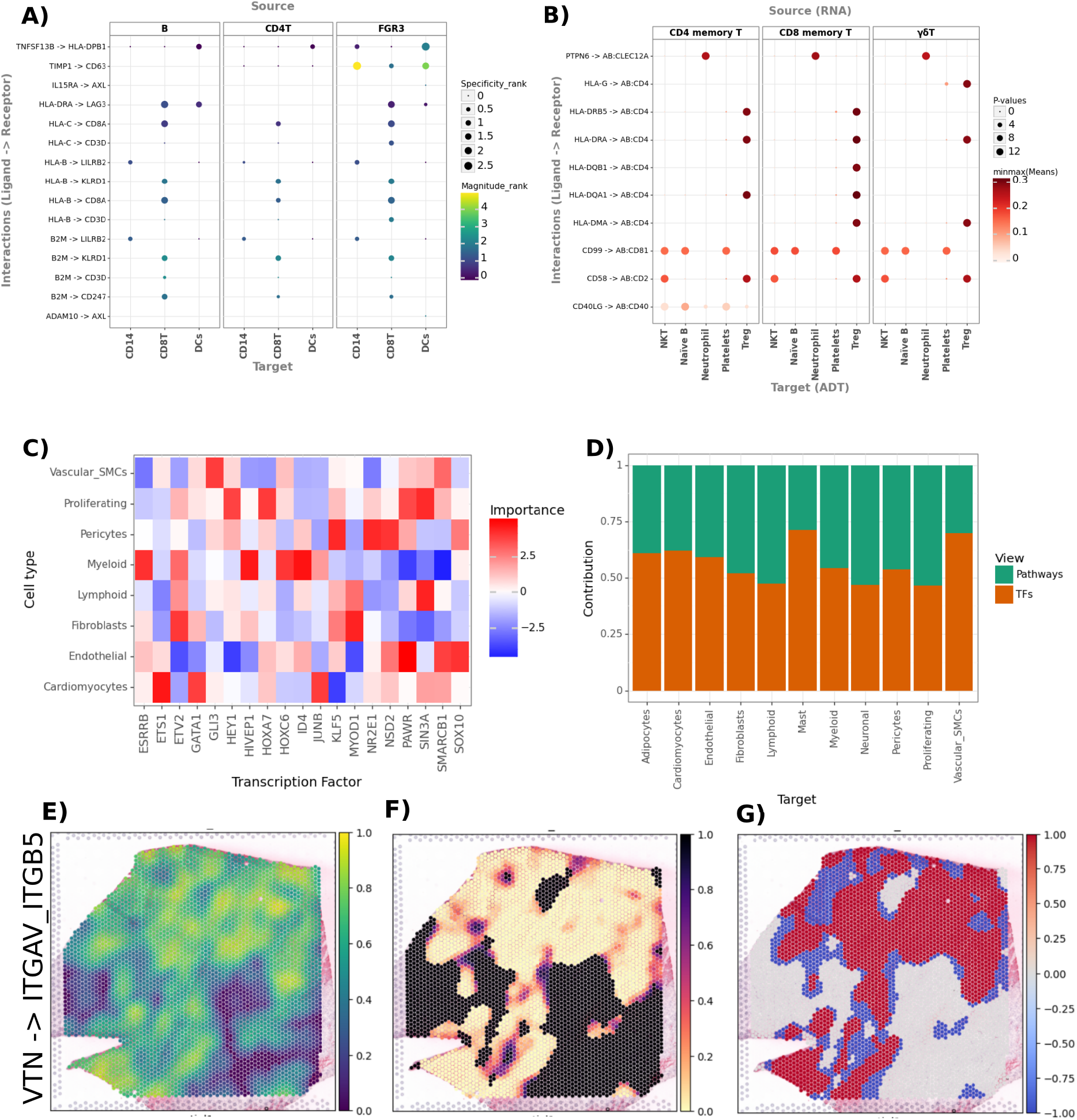
Inference of ligand-receptor interactions between peripheral blood mononucleated cells (PMBCs) from A) Lupus single-cell transcriptomics and B) Covid-19 CITE-seq (joined modality RNA-protein) data. C) Interaction importances (t-values) for transcription factors as predictors of cell type proportions and D) Contributions of distinct views (Pathways and Transcription Factors) as predictors of cell type proportions; in a 10X Visium slide of heart tissue with ischemia upon myocardial infarction. Local spatially-weighted E) Cosine similarity, F) Permutation P-values, and G) Categories for the interaction between VTN & ITGAV_ITGB5 in the ischemic heart slide from C and D.

LIANA+’s single-cell component is additionally applicable to multi-omics data (Methods). As an example, we adapted a permutation-based approach ^22^, initially proposed by CellPhoneDB ^3^, to the inference of ligand-receptor interactions from single-cell RNA and protein (CITE-seq) in human blood (Fig. 2B).

#### 2.1.2 LIANA+’s Spatial Component

As a consequence of the diverse array of spatial omics technologies and the varied tasks they encompass, a multitude of computational approaches are required to make most of the data. In this context, LIANA+ includes several strategies to flexibly analyse CCC from diverse spatially-resolved technologies (Fig. 1B).

First, we implemented global Moran’s R ^17^ - a bivariate extension of Moran’s I - used as a simple measure of spatial co-occurrence. Such co-occurrence measures, however, only consider two variables at a time, and hence do not account for complex relationships across variables. To this end, we also implemented a multivariate, multi-view modelling approach to learn spatial relationships across distinct types of features or spatial contexts (represented as views) ^16^. Our approach can learn complex relationships, such as relationships between ligand expressions and pathways ^16^, or cell types and pathways ^39,40^ (Fig. 2C), as well as jointly model any combination of views (Fig. 2D; Methods). As such, it is also well-suited to jointly learn relationships across combinations of data modalities and technologies. Thus, depending on the goal in mind, users might opt for a simple metric that summarises two variables at a time, or multi-view learning which not only accounts for collinearity between features, but also provides estimates about the predictive performance of different views.

The aforementioned methods infer global spatial relationships - i.e. they consider all spots to infer a value for each interaction across the slide as a whole. As such, they provide summary statistics for each interaction in a slide, but they do not provide information about the exact site or location of the interactions. To complement the global methods in LIANA+, we implemented six metrics to identify local interactions at the individual spot or cell location (Fig. 2E), along with permutation-based P-values for each (Fig. 2F; Methods). In brief, these are (i) an extension of univariate spatial clustering measures such as local Moran’s I (as implemented in SpatialDM ^17,41^); (ii) four spatially-weighted variants of commonly used metrics (Cosine similarity, Pearson and Spearman correlation, and Jaccard index); and (iii) a masked version of Spearman correlation, as proposed by scHOT ^18^ (Methods). Moreover, LIANA+ enables local interactions to be categorised according to the magnitude of expression or sign of the variables involved in the interaction (Fig. 2G; Methods).

To aid in the choice of local metrics, we evaluated their ability to preserve biological information in two tasks: (1) binary classification of malignant spots and non-malignant spots, and (2) cell type prediction, using for both tasks local ligand-receptor scores as predictors (Methods). Spatially-weighted Cosine similarity performed most consistently in both tasks (**Supp. Fig. S2**; **Supp. Note 1**). Thus, it was set as the default local metric in LIANA+, and also used for most analyses presented in this work.

In line with the single-cell component, the spatially-informed methods in LIANA+ are also easily applicable to multi-omics data. To demonstrate this, we analysed a recent, spatially-resolved tonsil CITE-seq dataset ^42^. Using our global multi-view learning approach, we found that the interaction between E-Cadherin protein expression and TP63 activity had the highest importance (OLS t-value = 7.96; Methods; **Supp. Fig. S3A-B**), both of which are associated with cell adhesion ^43,44^. To pinpoint the sub-region of the potential interaction, we used the local metrics, and saw high local scores predominantly within the boundary regions of vasculature (**Supp. Fig. S3C-F**).

Similarly, we used Global Moran’s R to identify co-clustered interactions between metabolite abundances, estimated from transcriptomics data (Methods), and their corresponding receptors. We saw that the interactions between the gamma-aminobutyric acid (GABA) neurotransmitter and several subunits of its corresponding GABA_A_ receptor had the highest, albeit relatively weak, global co-clustering; with subunit alpha 2 (Gabra2) having the strongest association to GABA (Global Moran’s R=0.18; **Supp. Fig. S4A-D**). Next, we employed a local metric to locate the brain region where the CCC event takes place. In line with previous observations ^45^, the interaction between GABA and Gabra2 predominantly potentially occurred within the cortex and hippocampus (**Supp. Fig. S4E**).

In conclusion, here we demonstrated that LIANA+ can be readily applied to identify spatially-informed interactions, driven by diverse mediators, and the corresponding regions in which they occur. We also showed how LIANA+ can combine existing components in new ways, as in the cases of spatial CITE-seq data and metabolite-mediated CCC.

#### 2.1.3 LIANA+ Multi-condition Component

As the number of samples and experimental design complexity in single-cell and spatial datasets continue to increase, generalizable methods are required to analyse CCC across conditions. In LIANA+, we provide several strategies to do so (Fig. 1C).

As an initial strategy and in line with current best practices of differential expression analysis in single-cell data ^46,47^, we used hypothesis-driven tests at the pseudobulk level with DESeq2 ^48,49^ to enable CCC inference between conditions (Methods). Briefly, LIANA+ combines differential analysis results with gene expression levels, and then aggregates those into ligand-receptor statistics across cell type pairs (Methods). This aggregation of feature-level statistics facilitates the prioritisation of potential communication events that distinguish groups of samples or conditions (Fig. 3A).

**Figure 3.**
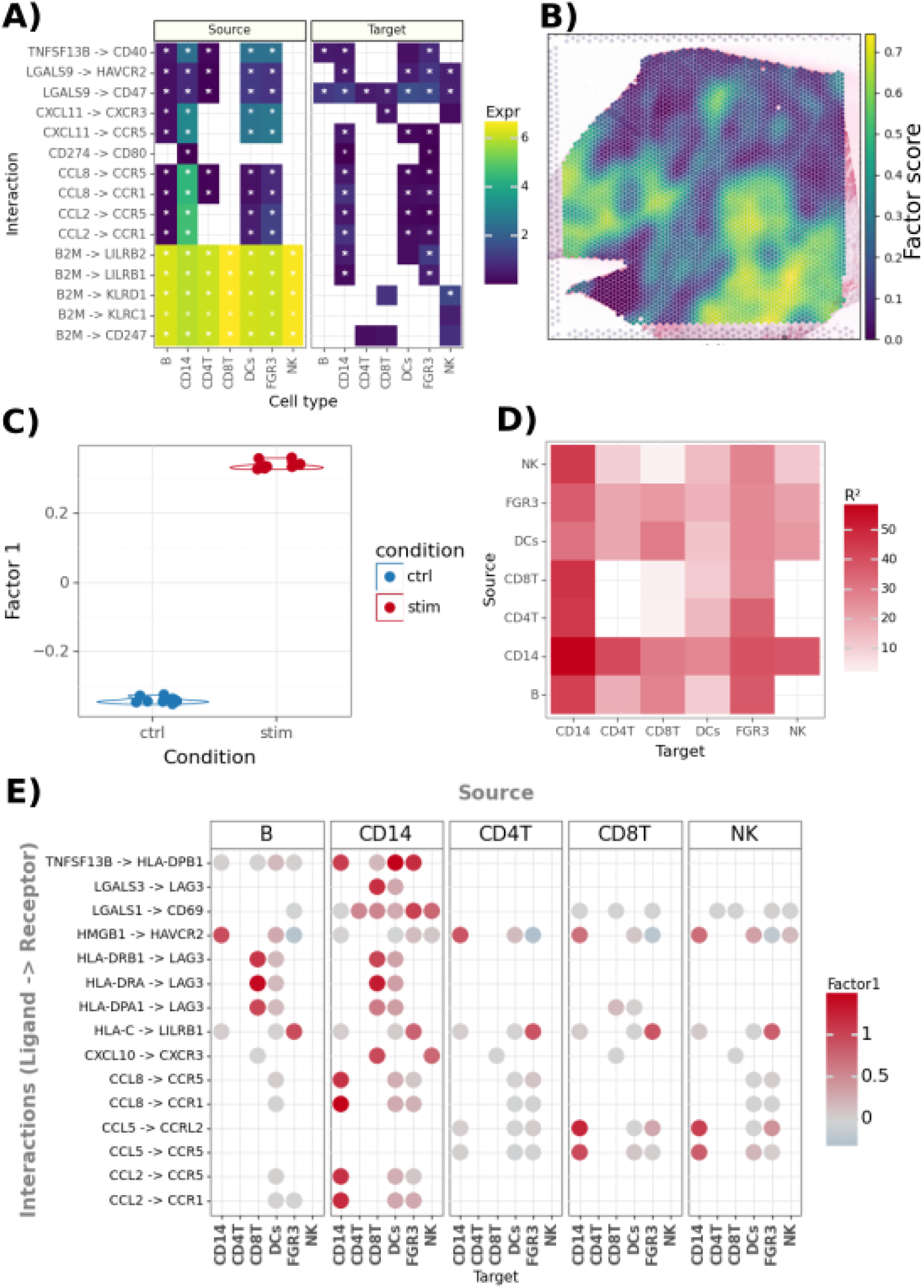
A) Hypothesis testing for deregulated ligand-receptor interactions upon interferon-beta treatment in lupus PBMCs (Methods). B) NMF factor scores on local ligand-receptor metrics in an ischemic heart slide. C) MOFA+ per-sample factor scores of decomposed ligand-receptor interactions from a factor (Factor 1) separating control and interferon-beta-stimulated lupus patient samples. Factor 1. D) Variance explained (R^2^) per cell type pair and interaction loadings in Factor 1.

CCC is a multicellular process, and while hypothesis testing for deregulation events of concerted pairs of ligand-receptor transcripts between two cell types can be helpful, such testing disregards coordinated communication events that involve multiple cell types. As an alternative, we combine CCC inference scores, both from dissociated and spatial data, with factorization approaches that allow for an unsupervised analysis of CCC of spatial locations or samples. These hypothesis-free (unsupervised) approaches utilise different dimensionality reduction algorithms to jointly model CCC events across samples and cell types, or locations. Specifically, they decompose inferred CCC events into latent factors that can be interpreted as intercellular programmes representing coordinated CCC events.

In spatial data, local spatially-informed metrics in LIANA+ can be combined with standard non-negative matrix factorization (NMF) (Fig. 3B), along with a heuristic approach to estimate the optimal number of factors (Methods). The application of NMF to local metrics results in intercellular programmes (factors), represented by factor scores per location, along with sets of interactions associated with each factor. In addition to being applicable to a single sample at a time, we also demonstrate that NMF can identify intercellular programmes in cross-condition data, as presented in Section 2.2.

For cross-conditional, dissociated single-cell data, LIANA+ leverages higher-order dimensionality reduction approaches to decompose CCC events into intercellular programs, as previously demonstrated with Tensor-cell2cell ^32,50^. Besides Tensor-cell2cell, we propose an alternative unsupervised approach that leverages the MOFA+ multi-view framework (Methods) ^51^. The approach inherits the efficiency and flexibility of MOFA+ to enable factor analysis of CCC interactions across samples, by modelling pairs of cell groups as views. In addition to providing information on factors that best separate the samples according to predefined conditions (Fig. 3C), both factorization methods also inform about the most relevant cell types (Fig. 3D) or interactions (Fig. 3E), explaining the variability of CCC events across samples.

In LIANA+, any ligand-receptor method can be combined with MOFA+ or Tensor-cell2cell. As such, we evaluated the ability of each combination of ligand-receptor method and dimensionality reduction to distinguish samples from different conditions (Methods; **Supp. Note 2**) using five public cross-conditional atlases from the human heart, lung, and brain (**Supp. Table 3**). We found that both Tensor-cell2cell and MOFA+ capture CCC events coordinated across cell-types that separate samples according to the different conditions at hand. Of note, while the results of most methods (except CellChat) were comparable when decomposed with MOFA+, there were more differences in method performance when using Tensor-cell2cell (**Supp. Fig. S5**; **Supp. Note 2**).

To showcase the combined approach of LIANA+ with MOFA+, we used public single-cell and spatially-resolved murine acute kidney injury datasets ^52,53^. In the dissociated dataset, we identified a potential intercellular programme modulating tissue repair in response to kidney injury (**Supp. Fig. S6A-D**; **Supp. Note 3**). We then used the spatial component of LIANA+ to confirm that an interaction involving Spp1, associated with kidney injury in dissociated data, was also captured in the spatial dataset. We found that the interaction increased in prevalence subsequent to injury (**Supp. Fig. S6E&F**), supporting its potential relevance.

#### 2.1.4 LIANA+ Intracellular Signalling Component

CCC events commonly initiate or emanate from intracellular processes, and LIANA+ provides various strategies to investigate these.

Leveraging OmniPath ^33^, LIANA+ enables the annotation of ligand-receptors to pathways of interest. It is also possible to perform downstream enrichment analyses of the output of any aforementioned CCC methods, from single-cell or spatial data, using a wide range of gene sets and enrichment methods via the package decoupler ^54^ (Fig. 4A; Methods).

**Figure 4.**
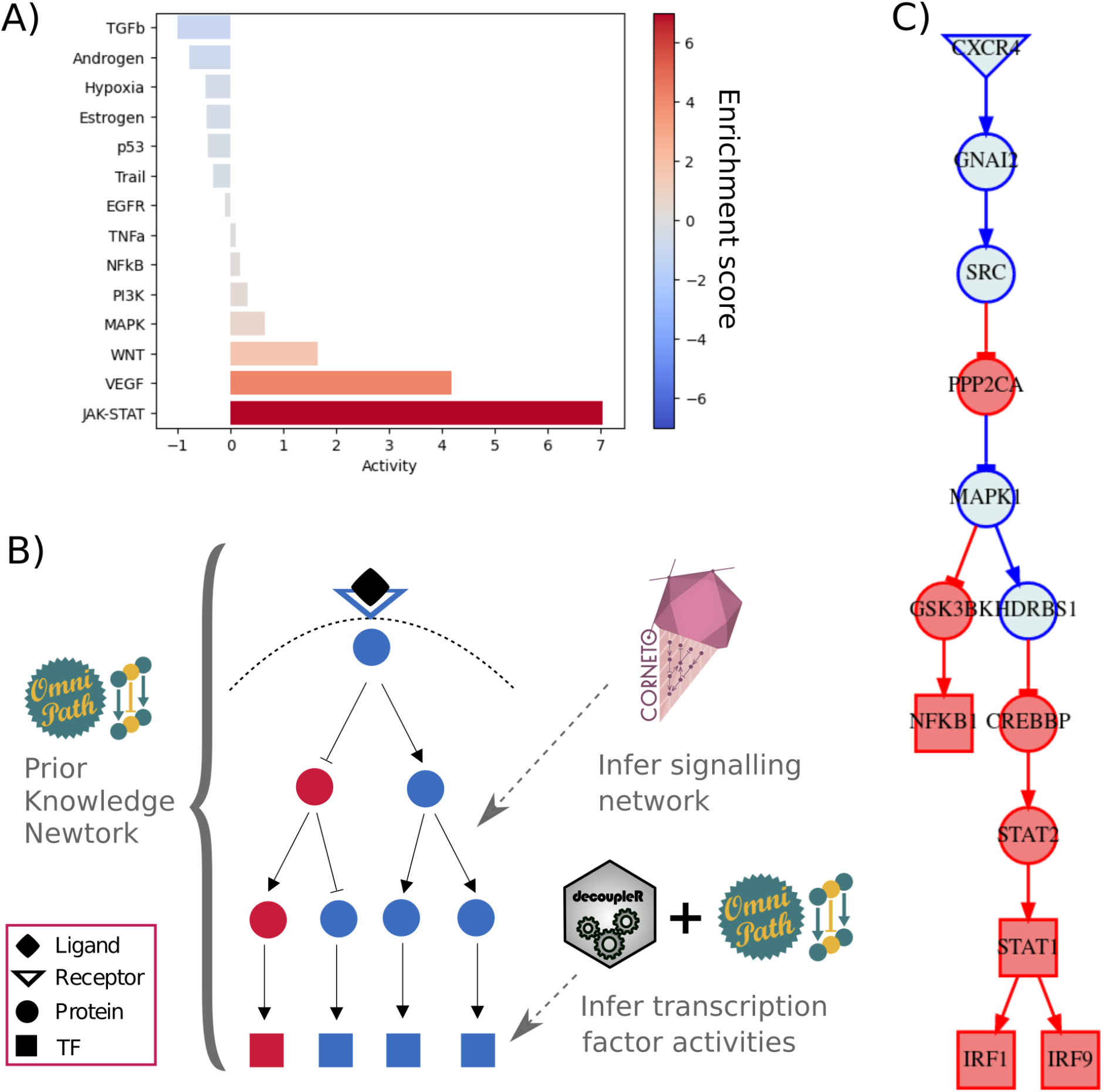
A) Pathway enrichment analysis of ligand-receptor loadings identified with Tensor-cell2cell, showing high JAK-STAT activity following interferon-beta treatment. B) Outline of workflow to link deregulated ligand-receptor to intracellular signalling. C) Causal intracellular signalling network, connecting deregulated CCC events following interferon-beta treatment with downstream transcription factors, associated with JAK-STAT signalling. All analyses in this figure were done using the lupus PBMCs dataset (Methods).

In addition, LIANA+ can infer signalling networks from prior knowledge, linking identified CCC events to downstream intracellular signalling pathways and transcription factors. In contrast to existing network methods used in the CCC field ^9,10^, our approach considers the direction of deregulation for nodes of interest (e.g. the activation or inhibition of receptors and transcription factors), as well as the signs of edges within the prior knowledge (activating or inhibiting edges; Methods). For this, CCC predictions from any method in LIANA+ are combined with knowledge of signed and directed protein-protein interactions as well as with transcription factors and their targets, both obtained from OmniPath (Fig. 4B; Methods). Then a network-optimisation approach ^55^ is used to identify putative causal paths that connect CCC events with active transcription factors (Fig. 4C; Methods). These analyses enable the user to obtain an integrated picture of the intra- and intercellular processes.

#### 2.1.4 LIANA+’s Prior Knowledge Component

All components of LIANA+ rely on existing biological knowledge. As such, LIANA+ draws from OmniPath’s rich database of ligand-receptor resources ^33^, providing access to 15 different resources, along with a consensus resource. To increase the flexibility of our CCC workflows, the knowledge in LIANA+ can be further expanded by leveraging BioCypher, which provides utilities for the modular and reproducible representation of knowledge ^56^. For instance, we created Metalinks ^57,58^ - a comprehensive and customisable resource of metabolite-protein interactions, additionally incorporating annotations such as tissues, pathways and diseases. We foresee that similar advancement will further refine the predictions from LIANA+.

### 2.2 LIANA+ extracts Condition-specific Communication Patterns

To showcase the ability of LIANA+ to decipher the intercellular mechanisms driving disease, we used a cross-condition spatial transcriptomics dataset (Visium 10x) ^59^ of human myocardial infarction encompassing samples from myogenic, fibrotic, and ischemic regions (Fig. 5A).

**Figure 5.**
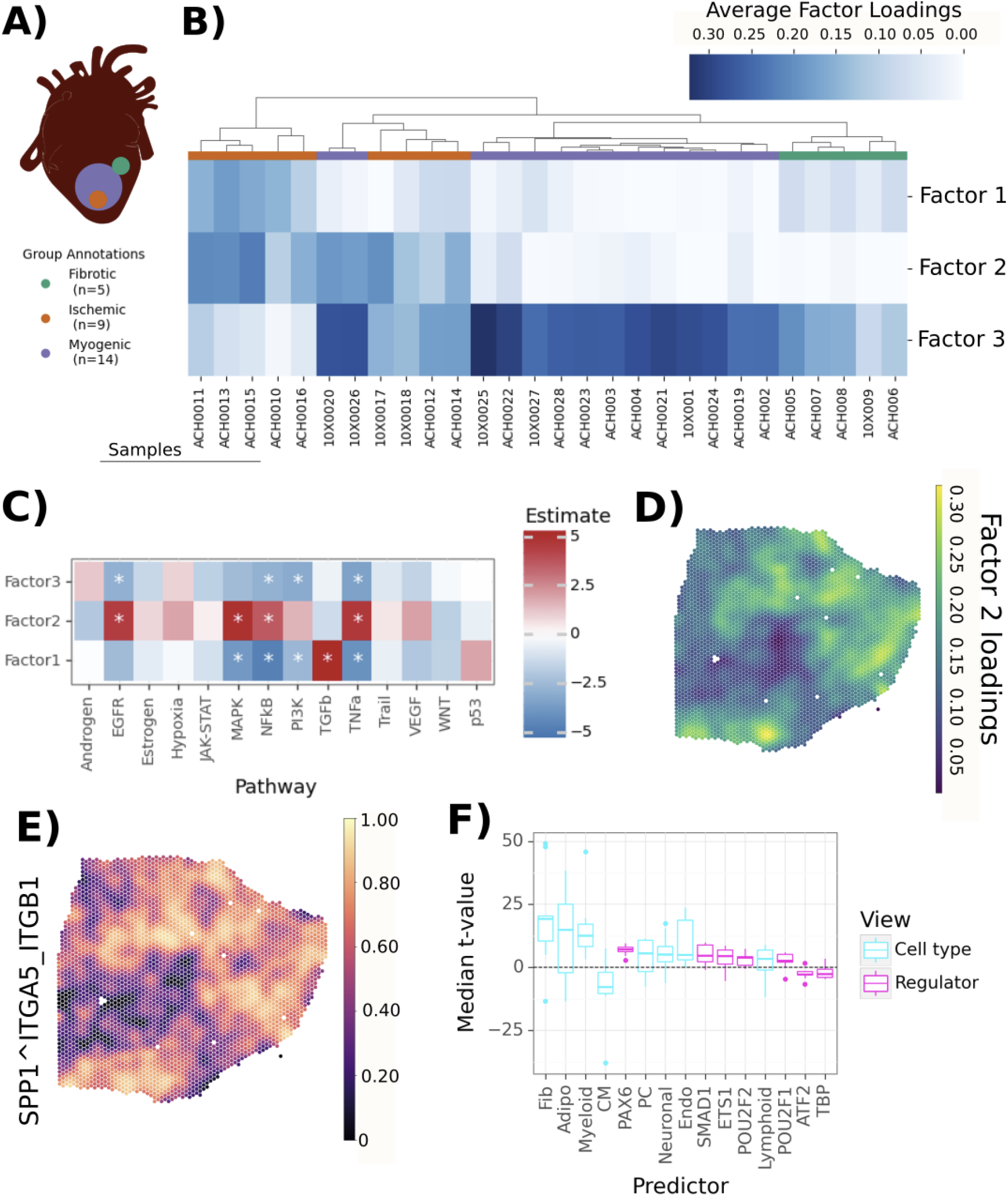
A) Experimental design. B) Average factor scores per slide. C) Pathway enrichment of ligand-receptor loadings. D) Factor 2 scores in a selected sample (ACH0014). E) Spatially-weighted Cosine similarity of SPP1 and the ITGA5_ITGB1 complex. F) Median importance (t-values) from the MISTy models for the local interaction between SPP1 and ITGA5_ITGB1; showing cell type proportions and transcription factors (regulators) as predictors in blue and pink, respectively.

To identify shared and coordinated CCC events across all slides, we used NMF on the local (spot-level) metrics of all ligand-receptor interactions, calculated using Cosine similarity. The “elbow” method approximated the presence of three factors as optimal (**Supp. Fig. S7A**; Methods), and we thus decomposed the local interactions into three factors, which we assumed to represent three distinct intercellular interaction programmes (patterns).

We then aggregated the factor scores for each slide, and saw that the slides clustered largely according to the regions from which they were obtained (Fig. 5B). We saw high mean scores of Factor 3 in Myogenic samples, while Factor 2 scores were prominent in Ischemic samples, and Factor 1 scores were high in both Ischemic and Fibrotic slides (Fig. 5B). To better understand the biological processes underlying the identified communication patterns, we did pathway enrichment analysis on the ligand-receptor loadings of each factor (Methods; Fig. 5C). The ischemia-associated Factor 2 showed an up-regulation of TNFα, NFkB, and MAPK pathways (Fig. 5C), reflecting expected inflammatory communication patterns in ischemic regions (Fig. 5D) ^60^. Similarly, we saw that interactions associated with TGFβ signalling, a well-known driver of fibrosis ^61^, were enriched in Factor 1.

To decipher the global drivers of ischemia-associated communication patterns, we modelled, in a spatially-informed manner, the top 20 most representative local ligand-receptor scores from Factor 2 using two distinct sets of predictors: tissue function, represented by transcription factor activities, and structure, quantified by the cell-type proportions in each location (Methods).

We saw that, across the slides, the top 20 local scores from the ischemia associated factor (Factor 2) were well explained by jointly modelling structure and function (multi-view median R^2^ >= 0.75; **Supp. Fig. S7B**), with transcriptional factors having a higher contribution to the predictive performance (>90%) than tissue structure (<10%; **Supp. Fig. S7C**). This can be explained by the broader signalling events captured by the higher number of predictors - in this case transcription factors. Among the top 20 interactions in Factor 2, we found several interactions involving SPP1, a well-known driver of fibrosis ^62^. Specifically we saw several interactions involving integrin complexes with high variances explained (R^2^ > 0.81; **Supp. Fig. S7B)**, and we focused on the interaction of SPP1 with the ITGA5_ITGB1 complex (Fig. 5E). We saw that the abundance of cell types showed the highest predictive performances of the local score of SPP1 and the ITGA5_ITGB1 integrin complex (Fig. 5F). In particular the presence of fibroblasts was the best predictor of this interaction (median coefficient t-value = 19.3), together with myeloid cells (median coefficient t-value = 12.6; Fig. 5F). It was previously reported from this data set that the location of SPP1+ macrophages is associated with the presence of myofibroblasts in ischemic heart tissue ^59^. Our expanded analysis revealed a collection of potential communication events that could be linked to the proliferation or establishment of the myofibroblast phenotype in myocardial infarction that had not been reported before. The transcription factor most strongly associated with the same interaction was PAX6 (median coefficient t-value = 6.8; Fig. 5F), with the functional perturbation of PAX6 being previously associated with myofibroblast differentiation ^63^. We examined the local spatial co-expression of SPP1&ITGA5_ITGB1 and its relationship with the PAX6 regulator, and saw that their local spatial association (**Supp. Fig. S7D-E**) was primarily within the boundaries of the ischemia-associated intercellular programme (Factor 2; Fig. 5C).

In summary, this analysis showcases how LIANA+ provides a suite of spatially-informed methods to enable the identification of disease-related communication patterns, as well as diverse strategies to decipher and interpret the drivers of underlying biological processes within those niches.

## 3. Discussion

As our ability to quantify molecular readouts at scale increases, so does the demand for comprehensive methods to generate biological insights. Building on existing methodological infrastructure ^36^ and biological knowledge ^33^, LIANA+ integrates and expands previous methodological developments to study CCC, redefining them into synergistic components.

Single-cell technologies capture cellular heterogeneity at an unprecedented scale, yet during the dissociation process information about tissue architecture is lost. Conversely, spatial technologies preserve tissue context, yet they either provide limited resolution since each spot captures multiple cells, or they measure a relatively low number of genes ^64^. LIANA+ addresses these limitations by combining CCC inference from single-cell and spatially-resolved data. In our application to murine kidney injury, we demonstrated how LIANA+ can identify interactions between cell groups of interest from single-cell data, and then support those by pinpointing corresponding spatially-informed interactions.

The components in LIANA+ are also complementary when applied to a single technology. For example, spatial methods in LIANA+ range from simple metrics, such as Global Moran’s R ^17^, to the spatially-informed modelling of multi-view representations ^16^. Both of these approaches summarise spatial relationships across the whole slide, and are thus highly complementary with local spatial metrics, six of which are implemented in LIANA+, to summarise interactions at the individual spot/cell level. As a consequence, LIANA+ provides ways to summarise CCC interactions relevant for the whole slide, but can also identify the specific subregions within which interactions occur.

Motivated by the rapid emergence of multi-omics, in particular spatially-resolved technologies ^23^, all methods in LIANA+ are also applicable across modalities. This was demonstrated in this work by jointly analysing RNA and protein omics layers from single-cell ^65^ and spatial ^42^ CITE-seq datasets. In our application to CITE-seq single-cell data, we showed that LIANA+ does not only adapt existing multi-omics CCC methods ^22^ but also provides readily usable novel strategies. As an example in the spatially-resolved CITE-seq dataset, we combined multi-view learning ^16^ with local metrics to learn spatial interactions across modalities and subsequently locate the subregion at which they occur.

Beyond analysing multimodal data, in LIANA+ we also augment conventional CCC inference by combining it with features derived from the data. For example, we built up on recent applications ^24,26,27,34^ to propose a strategy that estimates metabolite-mediated CCC, allowing the spatially-informed inference of interactions between neurotransmitters and their corresponding receptors. While this example uses only transcriptomics, the flexible nature of LIANA+ enables inferred metabolite abundance to be seamlessly replaced with quantitatively measured metabolite levels - to be enabled by emerging spatial metabolomics technologies ^66^, joined with transcriptomics.

These examples further underscore the ability of LIANA+ to be readily adapted to novel applications. Thus, we envision LIANA+ to be a versatile tool for the study of CCC driven by diverse mediators, beyond protein-mediated ligand-receptor interactions and expanding the range of CCC events that could be studied, such as host-microbiome interactions ^67–69^.

While each of the methods in LIANA+ are typically applied to a single slide or sample, we combine them with dimensionality reduction techniques to enable their application to multi-sample and multi-condition datasets. This enables the unsupervised analysis of coordinated CCC events as intercellular programmes, in line with approaches that aim to capture coordinated gene programs across cell types ^70–72^. Here, we repurposed MOFA+ ^51^ to CCC, and supported its ability to distinguish conditions in five cross-conditional single-cell atlases. We also used MOFA+ to identify intercellular programmes driving response to acute kidney injury in single-cell dissociated data. Moreover, we showed that factorisation techniques, such as standard NMF, can be combined with spatially-informed local interactions to identify cross-conditional programmes following myocardial infarction ^40^. While here we present each factorization technique in a specific application, they are interchangeable and in future applications, they can also be replaced with spatially-informed dimensionality reduction approaches ^73,74^. Moreover, such unsupervised approaches can be supplemented or replaced by a hypothesis-testing approach using differential expression analysis, as a simpler alternative that focuses on a single interaction at a time.

In addition to identifying CCC interactions relevant in both steady-state or across conditions, LIANA+ facilitates interpretation of intercellular signalling by connecting it to downstream events. Leveraging its flexibility, it can be integrated with diverse enrichment methods ^54^ and existing knowledge resources ^33^ to infer CCC-associated pathways or downstream transcription factors. LIANA+ includes a component ^55^ to infer causal paths connecting CCC to transcription factors. While here we used an algorithm to infer sign-consistent networks ^55^, other network approaches can also be incorporated ^6–10^. Consequently, LIANA+ enables the integration of both intra- and intercellular communication events.

The methods implemented in LIANA+ have a number of limitations. First, they use prior knowledge, which is limited, often exhibiting biases and a trade-off between coverage and quality ^9,11,75^. Most curation efforts have been focused on annotating ligand-receptor interactions ^3,4^, and additional prior knowledge efforts are needed in particular for the inference of CCC beyond protein-mediated events. Moreover, contextualising prior knowledge to specific cell types, tissues, or disease can help to reduce erroneous predictions. As an example, we made use of MetalinksDB ^58^, a comprehensive resource for the inference of metabolite-mediated CCC, here customised to the brain. Second, CCC from dissociated single-cell data remains limited to the co-expression of communication partners, and this co-expression at the transcript level may not translate to the protein level, let alone imply a functional interaction ^11^. Likewise, while spatially-resolved data is a step further from its dissociated counterparts, it is limited to the co-localization of transcripts. Finally, while we showed the ability of LIANA+ to generate CCC insights across a range of technologies, along with some preliminary evaluations, systematic benchmarks of CCC methods are still pending. Some examples exist but they remain limited to the use of orthogonal modalities such as spatial data ^11,76^ or downstream signalling ^11^. As emerging technologies ^77–79^ which provide *bona fide* CCC events, become measurable at scale and widely available, LIANA+ will serve as a facilitator for such benchmarks and comparisons. Therefore, despite its broad functionalities, our framework - as all CCC methods - remains a tool for hypothesis generation, requiring validation experiments.

Overall, LIANA+ generalises the multifaceted aspects of cell-cell communication inference into synergistic components. As illustrated in this manuscript, these components can be combined in various ways, and their configurations can be tailored to address diverse questions and datasets. Given the modularity of LIANA+, new methods can be integrated into the framework and immediately tap into the established ecosystem of methods and resources, benefiting from enhanced compatibility and interoperability. Thus, LIANA+ not only stands as a comprehensive and scalable tool for studying communication events but also serves as a foundational framework and catalyst for future collaborative developments in the field.

## 4. Methods

### Bivariate Spatially-informed Metrics

Common notations:

*x_i_* and *y_i_* are the vectors of two variables for spots (or cells) *i*

*n* is the number of spots,

*x* and *y* is the mean of the variable values,

*w_ij_* is a spatial proximity weight indicating the degree of spatial association between spot *i* and spot *j*.

We adapted bivariate Global and local Moran’s R, extensions of Moran’s I ^41^, from SpatialDM ^17^; both of which are measures of spatial co-occurrence.

Local Moran’s R is defined as:

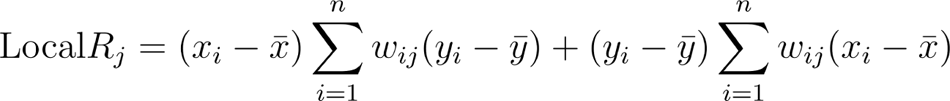

while Global Moran’s R is defined as:

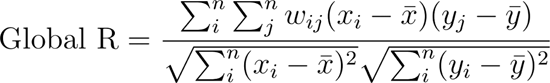

Inspired by scHOT ^18^, we also implemented local weighted variants of common similarity metrics, such as Pearson and Spearman correlation:

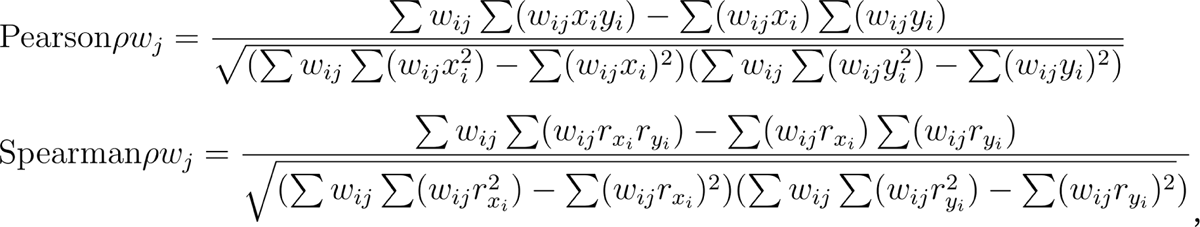

where summation is performed over i and *r_xi_*, are the ranks of *x* and *y* for spots *i*.

A second masked version of Spearman correlation as proposed and default approach in scHOT was also implemented; where we consider *r_xi_, r_yi_* only for spots *i* with non-zero *w*.

Moreover, we provide weighted Jaccard and Cosine similarity metrics:

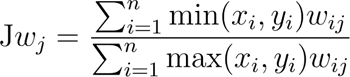

where:
*x_i_* and *y_i_* are vectors of the same length, binarized by setting values > 0 to 1, to signify presence or absence of a read out.

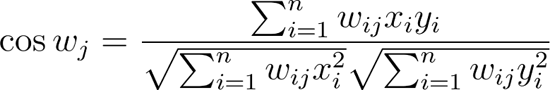

When working interactions, the members of which contain heteromeric complexes, we consider the minimum expression of complex members per spot. Any interactions, the members of which are not expressed in at least 10% of the spots are excluded.

### Local Score Categories

Inspired by GeoDa ^80^, we categorise local bivariate associations according to the magnitude and sign of the two variables. If spatially-weighted variables are non-negative (e.g. gene expression) then they are z-transformed. Then for each spot *j*, we categorise interactions according to the sign of the spatially-weighted variables (*v*) involved in the interaction - i.e. as positive, negative or neither:

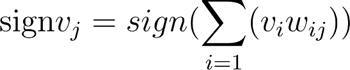

Then to obtain a category for the interaction, we combine the sign of the two variables (*x* and *y*). If both variables are positive (high-high), then the interaction is positive (1); if one variable is positive (high) and the other negative (low) then the interaction is negative (−1); if both variables are negative (low-low), or either variable is neither (e.g. equals to 0), then the interaction is labelled as “neither” (0). The latter enables us to distinguish relationships where both variables are highly-abundant (high-high) from those where both values are lowly-abundant (low-low).

For statistical testing of local metrics, we use spot label permutations to generate a Null distribution against which empirical local p-values are computed.

We provide a detailed tutorial on the bivariate metrics at: https://liana-py.readthedocs.io/en/latest/notebooks/bivariate.html

### Learning Spatial Relationships with MISTy

To learn multivariate interactions in space that go beyond bivariate metrics, we re-implemented MISTy’s multi-view learning approach ^16^. Our multi-view approach jointly models different spatial and functional aspects of the data, such that it can fit any number of views, and each view can contain any number of variables. As shown in this work, one can use it to jointly model different combinations of RNA expression, cell type proportions, pathway or transcription factor activities, CCC events, and protein abundance.

We additionally make use of different modelling approaches, by default, the models are based on random forests (RF) and can capture complex non-linear relationships. Here, we also implemented linear models. For both RF and linear models, we use the implementations available from scikit-learn ^81^.

In LIANA+, multi-views are represented as subclasses of MuData ^82^, modified to ensure the correct format of the views and corresponding spatial connectivities. Each multi-view structure has an intrinsic view (intraview) that contains the target variables of interest for each spatial location. The other views can be considered as “extra” views, composed solely of predictor variables. Predictor variables can also represent a transformation of the variables within the intraview taking into account a specific spatial context, as well as other categories of variables.

Once the multi-view structure is defined, each target is modelled by predictors from each view independently. As such, for each target we obtain (1) relationship importances for each of the predictors from the distinct views; (2) the relative ‘contribution’ of each view to the joint prediction of each target (3); as well as the goodness of fit (e.g. R^2^) of the model.

The way that importances of target-predictor relationships (1) are calculated depends on the modelling approach. For RFs, we use the reduction of variance explained that can be attributed to each predictor across all regression trees. For the linear models we use the t-statistic of the estimated parameters under the zero value null hypothesis. The independent view-specific predictions are combined by a cross-validated regularised linear meta-model ^16^ to obtain the contributions of view-specific models (2), along with the goodness of fit of the overall model, for each target variable (3). In particular, we can discern between the contribution of the intraview, modelled as the intrinsic variability among target variables within the same cells/spots, from the predictive contribution of “extra” views, which encode spatial information.

To facilitate the use of our multi-view learning approach, we provide in depth tutorials on how to generate and model custom and predefined multi-view structures: https://liana-py.readthedocs.io/en/latest/notebooks/misty.html.

### Estimation of Spatial Connectivities

As in MISTy ^16^, spatial connectivity weights are calculated using families of radial basis: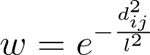, Gaussian 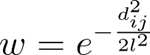, linear 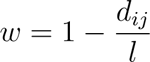, and exponential kernels 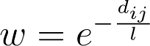; where *w* is a weight matrix of shape *n* x *n*, *d_ij_* is the Euclidean distance between cells or spots *i* and *j*, *l* is a parameter controlling the shape, or bandwidth. We additionally provide a cutoff parameter below which spatial connectivities are set to zero. Throughout the manuscript, unless otherwise specified, we used Gaussian weights with a bandwidth of 150, and a cutoff of 0.05, and the diagonal (self to self) was set to 1 for local scores and 0 for MISTy.

Programmatically the calculation of spatial connectivities mirrors Squidpy’s `spatial_neighbors` function, and thus spatial connectivities can be easily replaced with Squidpy’s neighbourhood graphs ^83^.

### Ligand-Receptor Pathway Enrichment

To perform ligand-receptor pathway enrichment, we first convert gene set (pathway) resources, represented as weighted bipartite graphs where each gene belongs to a gene set, into ligand-receptor sets. Specifically, we assign a weight to each ligand-receptor interaction, based on the mean weight of the ligands and receptors involved in the interaction, also taking into account the presence of heteromeric subunits. Moreover, we assign a given ligand-receptor interaction to a specific gene set (or pathway), only if all members of the interaction are part of the gene set, and in the case of weighted resources are additionally sign-consistent. Finally, once a ligand-receptor resource is generated, we use decoupler-py to perform enrichment with univariate linear regression ^54^.

In this manuscript, we used the PROGENy resource ^84^ to assign pathway annotations to ligand-receptor interactions. In contrast to classic pathway gene sets, PROGENy contains consensually regulated targets of pathway perturbations ^84^, not genes thought to be members of the pathways. However, this resource-conversion procedure is applicable to any resource, including undirected resources, such as GO terms for which all members of a gene set will be assigned a weight of 1.

### Hypothesis testing for deregulated CCC across Conditions

To enable hypothesis testing for CCC, similarly to the strategy in MultiNicheNet ^85^, we first generate pseudobulk profiles by summing raw expression counts for each sample and cell type with the decoupler-py package ^54^. After filtering low quality genes in (e.g. considering minimum expression in terms of total counts and samples in which the gene is expressed), we perform differential analysis for each cell type independently with DESeq2^48^, as implemented in PyDESeq2 ^49^.

Once feature statistics per cell type are generated, we transform those into a dataframe of interaction statistics by joining them to a selected ligand-receptor resource, while additionally calculating average feature expression and expression proportions per cell type, based on a user-provided AnnData object^86^. Similarly to any other method in LIANA+, interactions expressed in less than 10% (by default) of the cells per cell type are filtered, considering the individual members of heteromeric complexes.

A detailed tutorial is available at: https://liana-py.readthedocs.io/en/latest/notebooks/targeted.html

### Sign-consistent Intracellular Networks

By combining CCC predictions with prior knowledge networks of intracellular signalling, it is possible to recover putative causal networks linking CCC events to transcription factors. To accomplish this, we used CORNETO ^87^ - a Python package that unifies network inference problems from prior knowledge - to implement a modified version of the integer linear programming (ILP) formulation implemented in CARNIVAL ^88^.

This modified version of CARNIVAL ^88^ takes four distinct inputs: (1) a prior knowledge graph (PKN) of signed protein-protein interactions, where nodes are proteins and edges are activating or inhibitory interactions; continuous and signed (2) starting (input) nodes and (3) end (output) node values, with negative values indicating downregulation and positive values indicating upregulation. In addition, we take (4) values for the rest of the nodes in the graph [0, 1] (e.g. gene expression proportions), with higher values incurring less penalty than genes with lower values when the gene is included in the inferred network. Then, a subnetwork, optimised for sparsity, is extracted from the PKN which connects the input (starting) nodes to the output (end) nodes.

The resulting inferred network is a directed acyclic graph that connects the (2) input nodes to the (3) end nodes (e.g. receptor to transcription factors), including the values for each edge and node of the graph indicating if the node is upregulated (+1), or downregulated (−1). A node n_c_ in the graph can be upregulated only if there is at least one selected parent node n_p_ such that n_p_ is upregulated and there is an activating edge between n_p_ and n_c_, or n_p_ is inhibited and there is an inhibitory edge between n_p_ and n_c_. Similarly, a node n_c_ can be downregulated if there is a parent node n_p_ downregulated with an activating edge between n_p_ and n_c_, or if a parent node n_p_ is upregulated and there is an inhibitory edge between n_p_ and n_c_.

These rules are encoded using linear constraints and continuous/binary variables to define a Mixed ILP problem, which is a particular type of a combinatorial problem with linear constraints. The optimization problem is defined as:

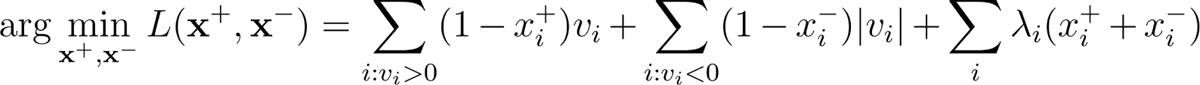

where is a vector of binary variables for each node in the PKN indicating whether the node *i* is upregulated (*x^+^_i_ = 1*) or not; *x^−^* is a vector of binary variables for each node *i* in the PKN indicating whether node in the PKN is downregulated (*x^−^_i_ = 1*) or not; *v* is a vector of values for measured nodes (input nodes and output nodes), where positive values are upregulated species and negative values are downregulated species. For example, *v* can be estimated as fold change, t-statistic or any other score indicating difference in activity in a protein in the PKN between two conditions.;

In this modified version of CARNIVAL, we additionally introduce λ - a vector of penalties to penalise the inclusion of protein nodes in the resulting inferred network, according to (4) node weights [0, 1] in the (1) PKN.

We set λ to penalty_max_ (1 as default) and penalty_min_ (0.01 as default):

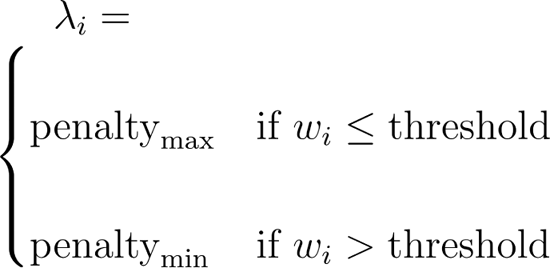

Linear constraints impose conditions on the variables of the ILP problem. For example, a node cannot be upregulated and downregulated at the same time (*x^+^ + x^−^ ≤ 1*). The problem includes other variables and linear constraints to guarantee that the final networks valid solutions are acyclic networks and that the rules explained before are respected. Additional information about the formulation can be found in Liu et al. (2019) ^55^.

We show the inference of sign-consistent networks downstream of deregulated CCC events, identified using differential expression analysis with PyDESeq2 ^49^ here: https://liana-py.readthedocs.io/en/latest/notebooks/targeted.html

### NMF on ligand-receptor local scores

A utility function was implemented that takes an AnnData object ^86^ as input and uses Scikit-learn’s NMF implementation to factorise the input matrix into two matrices of dimensions *k*, *n* and *k*, *m*; where *m* is the number of features, *n* is the number of observations (cells); and *k* is the number of components (factors). To estimate k, we additionally provide an heuristic elbow selection procedure, in which the optimal component number (*k*) is chosen from a sequential range of components using elbow selection as implemented in the kneedle package ^89^. Selection of optimal *k* is based on the mean absolute reconstruction error.

### LIANA+ in multimodal single-cell & spatial data

To enable the inference of CCC across modalities, the methods implemented in LIANA+ accept MuData objects ^82^ as input. These essentially provide functionalities to load and store multimodal data ^82^, and can be thought of as an extension of AnnData objects ^86^, which are the default input of LIANA+ when working with unimodal single-cell or spatial data.

Feature level transformations such as z-scoring or min-max scaling are used to transform features within omics and across omics to a comparable scale to facilitate integration.

### Intercellular Communication Factorization with MOFA+

Inspired by the CCC factorization approach proposed in Tensor-cell2cell ^32^ and building on our recent application of MOFA+ to dissociated cross-condition atlases ^71^, we use the ligand-receptor inference methods from LIANA+ across each sample independently, and then transform this into a multi-view structure of cell type pairs (views), each represented by samples and the ligand-receptor interaction scores in each. To build the multi-view structure, we use the MuData format ^82^, and only views with at least 20 (by default) interactions in at least 3 (by default) samples are kept. Moreover, we exclude samples if they have less than 10 interactions (by default) and interactions are considered only if they are present in at least 50% of the samples (by default). Then we use the MOFA+ statistical framework ^51^ to decompose the variance of the samples into intercellular communication programmes.

A tutorial on extracting intercellular programmes from single-cell dissociated data with MOFA+ is available at: https://liana-py.readthedocs.io/en/latest/notebooks/mofatalk.html

### Spot Calling using Local metrics

To benchmark how well each local score in LIANA+ preserves biological information, we devised spot classification and regression tasks. In the spot classification task, we used four public breast cancer 10X Visium slides ^90^, with annotations labelled as malignant (containing “cancer” in their annotation) or non-malignant spots (any other spot). For each slide, we calculate local ligand-receptor scores using the local metrics in LIANA+. Then for each local metric, we trained and evaluated Random Forest Classifiers, with 100 estimators, using a Stratified K-Fold cross-validation strategy (k=10). AUROC and weighted F1 were calculated on the test sets, and their average across the folds was used in visualisations.

In the regression task, we used a public dataset with 28 10x Visium slides from left-ventricle heart tissues to compare how well different local metrics capture cell-type specific ligand-receptor events. In particular, we checked how well do the local scores LIANA+ predict cell type proportions per spot, inferred using cell2location ^91^ as done in Kuppe et al., 2022 ^40^. We used a Random Forest Regressor, with 100 estimators, utilising a K-fold cross-validation training strategy (k=5), and calculated the variance explained (R^2^) and root mean squared error for each score. All classification and regression tasks were performed through Scikit-learn (v.1.10.1).

For the inference of ligand-receptor interactions throughout this work, we used LIANA’s consensus resource - a resource combining the curated ligand-receptor resources in OmniPath ^33^.

### Sample Label Classification

For the condition classification task, building on a similar approach ^32^, we used public, pre-processed, cross-conditional atlases (**Supp. Table 3**), each selected such that they include more than five samples per condition following preprocessing. To ensure that only high-quality samples were used in each of the atlases, we removed any samples with less than 1,000 cells or z-transformed total counts above or below a z-score of 3 and −2, respectively. In the Carraro et al. dataset ^92^, we kept samples with more than 700 cells. Moreover, only cell types found in at least 5 samples and with at least 20 cells in each individual sample were considered. To ensure that the samples were balanced between the conditions, if either condition had a sample ratio higher than 1.5 x the number of samples in the other condition, then the overrepresented condition was subsampled to the number of samples in the underrepresented one. Each dataset was normalised to 10,000 total counts per cell and log1p-transformed.

Subsequent to preprocessing, we inferred ligand-receptor interactions at the cell-type level using the homogenised methods in LIANA+, independently for each sample. Any interactions not expressed in at least 10% of the cells in both source and receiver cell types were filtered.

Then the output from LIANA+ was converted to the structures used by the factorization approaches employed by MOFA+ and Tensor-cell2cell - a multi-view and a 4D tensor, respectively. For the factorization in both, we consider interactions only if they were present in 33% of the samples, and any interactions missing in a sample were assumed to be biologically-meaningful and assigned as zero. For all datasets, we decomposed the CCC events into 10 factors, except Reichart et al ^93^, which was decomposed into 20 factors due to its larger sample size.

Using the factor scores for each method-factorization approach combination we then performed a classification task, modelled to the one from Armingol et al. ^32^. Specifically, a Random Forest Classifier, with 100 estimators, was trained and evaluated on the sample factor scores computed for each score-factorization combination, utilising a Stratified K-Folds cross-validation strategy (k=3), performed over 5 seeds. Then the mean Area Under the Receiver-Operator Curve (AUROC) and weighted F1 scores were then calculated on testing set’s the probabilities and label predictions, respectively.

### Analysis of Single-Cell Data from Lupus Patients

We used a pre-processed dataset of 8 pooled patient lupus samples ^94^, each before and after interferon-beta stimulation. An AnnData object with the processed data was obtained from https://figshare.com/ndownloader/files/34464122; available via pertpy ^95^. Raw gene counts were normalised to 10,000 total counts per cell and log1p-transformed.

For the factorisation of CCC events with MOFA+, we used LIANA’s consensus rank aggregate of magnitude ligand-receptor scores (**Supp. Table 2**), focusing on ligand-receptors with at least 20 interactions in 30% of the samples, and views with at least 10 samples. Missing interaction values were filled with zeroes. Factorisation was also carried out with Tensor-cell2cell using the consensus rank aggregate. A 4D tensor was built such that all cell types were preserved, and Tensor-cell2cell estimated an optimal rank of six components. Ligand-receptor pathway enrichment analysis was then performed on the interaction loadings using multivariate linear regression ^54^, with gene annotations from PROGENy ^84^. For hypothesis-testing with PyDeSeq2, for each pseudobulk profile we considered only genes with counts in at least 5 of the samples, and at least 10 counts across all samples. Differential testing was performed between stimulated and control samples, using control as the reference. Then we used only the samples subsequent to interferon-beta stimulation to calculate gene expression and proportion statistics. We kept only interactions, all members of which were expressed in at least 10% of the cells in both source and target cell types.

For the inference of the downstream signalling events, we obtained a protein-protein interactions resource from OmniPath, considering only interactions with consensus direction, and a curation effort >= 5. We estimated the transcription factor activity using the Wald statistics from PyDeSeq2 with univariate linear regression ^54^ and CollecTRI ^96^. Then using CORNETO (v0.9.1-alpha.3), we inferred causal networks between receptors from the top 10 interactions, in terms of mean interaction Wald statistic, and the top 5 transcription factors, with highest enrichment scores, in CD14 monocytes. We used gene proportions with a cutoff of 0.1, such that nodes above the cutoff were assigned a penalty of 1, and those below a penalty of 0.01. An edge penalty of 0.01 was also used, and the problem was solved with the HIGHs solver ^97^, available within SCIPY (v1.10.1) ^98^.

### Analysis of Dissociated Single-cell CITE-seq Data

We obtained a single-cell CITE-seq dataset of peripheral blood mononucleated cells from COVID-19 positive controls ^65^. We normalised the gene expression and ADT counts using log1p and centred log-ratio normalisation, respectively. Then we inferred ligand-receptor interactions, using the normalised gene expression for the ligands and protein abundances for receptors. Both assays were feature-wise transformed using zero-inflated minmax, prior to applying an approach similar to CellPhoneDBv2 ^22^.

### Analysis of Spatial CITE-seq data in human Tonsil

We obtained a processed secondary lymphoid (tonsil) tissue with 273 measured proteins captured via barcoded antibodies and genome-wide gene expression ^42^. For transcriptome counts we used total count and log1p normalisation, while for protein abundances we used centred log-ratio normalisation. We then estimated transcription factor activities as a way to reduce the dimensionality and improve the signal within the dataset. Decoupler ^54^ was used to estimate the transcription factor activity, based on univariate linear regression, with DoRothEA ^99^. We then used our multi-view learning approach ^16^ to explore the spatial relationships between proteins and transcription factor activities, modelling predictions and importances for individual views with multivariate linear regression. We used feature-wise zero-inflated minmax transformation ^22^, with values below 0.25 being set to 0, to calculate spatially-weighted Cosine similarity.

### Analysis of Murine Acute Kidney Injury

We first filtered the preprocessed single-cell dataset, with pre-annotated cell types, to only those cell types with at least 15 cells in at least 10 samples; additionally excluding urothelial cells as they are not expected to communicate with most of the other cell types in the kidney. Following total count and log1p normalisation, we inferred ligand-receptor interactions using LIANA’s consensus method (**Supp. Table 2**), excluding any interactions not expressed in at least 10% of the cells in both the source and target cell types. Then we transformed the resulting ligand-receptor interactions into views representing cell type pairs, keeping only those interactions present in at least 25% of the samples, views with at least 15 interactions and at least 5 views. Finally, we decomposed those views into 5 factors using MOFA+. Kruskal Wallis test was performed on the sample loadings for Factor 1.

For the spatial data, we filtered the preprocessed 10x slides, such that only spots with at least 400 genes expressed and genes expressed in at least 5 spots were kept. We additionally excluded any spot outliers according to mitochondrial, ribosomal and total count content, using comparable but slide-specific thresholds. Then for the interactions of interest identified in the dissociated datasets, we calculated local cosine similarity and global Moran’s R.

### Analysis of Human Myocardial Infarction

We first estimated ligand-receptor local scores using Cosine similarity independently on each of the processed 10X Visium transcriptomics slides. We considered interactions whose members were expressed in at least 10% of the spots. Then we concatenated the resulting ligand-receptor AnnData objects (slides), and kept only those interactions present in at least 10 of the slides. Subsequently, we factorised the concatenated object with NMF, calculated the average factor scores per slide, and hierarchically clustered the results using Euclidean distances.

Pathway activities of ligand-receptor interaction loadings were calculated using linear regression ^54^ and sets of ligand-receptor pathways, annotated using the PROGENy pathway resource ^84^ with all genes.

Focusing on the top 20 interactions, in terms of highest loadings, from ischemia-associated (Factor 2), we modelled a multi-view representation such that the interactions were treated as targetes (intra-view), while cell type proportions and pathway activities were two distinct extra (predictor) views. Transcription factor activities were calculated using linear regression with the DoRothEA resource ^99^, while cell type proportions were inferred with cell2location ^91^ and available in the processed slides ^40^. We excluded proliferating cell type proportions when modelling because their phenotype is independent of cell-type lineages. This process was done for each slide from ischemic heart regions and aggregated statistics, such as ordinary least squares t-values, contribution and goodness of fit, were calculated using the median.

An ischemic slide (accession number: ACH0013) from this dataset was also used in the examples presented throughout the manuscript.

### Analysis of Metabolite-mediated CCC in Spatial Mouse Brain Data

In line with recent developments, we used simple enrichment-like approaches to estimate the abundance of metabolites using the expression of their corresponding enzymes ^24,26^. To generate the required prior knowledge, we made use of MetalinksDB ^100^ - a comprehensive metabolite-protein knowledge graph (KG). Briefly, MetalinksDB is built from metabolite-protein interactions extracted from databases ^26,27,101^ and genome-scale metabolic models ^102,103^ as an annotated KG using BioCypher adapters ^56^. This KG was customised to only include metabolites found in the brain or cerebrospinal fluid. Using this customised KG we generated: (1) a consensus resource of manually curated metabolite-receptor interactions; (2) sets of producing and degrading enzymes, respectively weighted as 1 and −1; (3) sets of transporters for each metabolite, with exporters being assigned to 1 and importers −1.

We then use a univariate linear regression model ^54^ to estimate metabolite abundances for each cell/spot. In a second step, inspired by NeuronChat ^26^, we calculate a transporter (export) score for each metabolite using a simple arithmetic mean, such that estimated metabolite abundances in each cell/spot, the export score of which is negative or 0, are set to 0. The metabolite estimation step in LIANA+ can be replaced by other more informed models ^104^, at the user’s discretion.

We used the approach, described above, to estimate metabolite abundances from a preprocessed adult mouse brain 10x Visium slide. Then we calculated Global Moran’s R and local Cosine similarity on feature-wise minmax-transformed metabolite abundances and normalised receptor gene expression. We additionally calculated P-values for the local Cosine similarities using 100 permutations.

## Data Availability

All datasets used in this work are publicly available.

## Code Availability

The latest version of LIANA+ is available at https://github.com/saezlab/liana-py, along with detailed tutorials describing the distinct components presented here (https://liana-py.readthedocs.io). LIANA+ is regularly released on Github and stable versions are released on PyPI (https://pypi.org/project/liana/).

## Acknowledgements

D.D. is supported by the European Union’s Horizon 2020 research and innovation program (860329 Marie-Curie ITN “STRATEGY-CKD”). S.L has received funding from the European Union’s Horizon 2020 research and innovation programme under grant agreement No 965193 (DECIDER). P.S.L.S. has received funding from the Deutsche Forschungsgemeinschaft under grant agreement SPP 2395. P.R.M. is supported by the European Union’s Horizon 2020 research and innovation programme under grant agreement No 951773 (PerMedCoE). R.O.R.F. is supported by DFG through CRC/SFB 1550 “Molecular Circuits of Heart Disease”.

We further thank Alberto Valdeolivas, Pau Badia i Mompel, Olga Ivanova, Martin Garrido-Rodriguez, Haikuo Li, Benjamin Humphreys, Nathan Steenbuck, Gregor Sturm, and Reyes Becerra Perez for the helpful discussions. We also thank Sophia Müller-Dott for her help and critical input to the graphical abstract.

## 6. Authors contributions

D.D. and J.S.R. conceived the project. D.D. wrote the manuscript and developed the software with input from other authors. The manuscript was drafted by D.D., edited by J.S.R. and jointly revised by all co-authors. All co-authors contributed to the case studies and writing and gave feedback for software development and/or text, coordinated and integrated by D.D. J.S.R supervised the work. All authors approved the final manuscript.

## Appendix

### Supplementary Note 1. Spot calling Evaluation

First, we evaluated the ability of the spatially-weighted local metrics to classify malignant and non-malignant spots in four breast cancer slides ^90^ (Methods). All scoring functions did well at classifying malignancy (AUROC > 0.9; weighted F1 > 0.85; **Supp. Fig. S2A-B**). Spatially-informed Jaccard and Cosine similarity functions had slightly higher and more AUROCs and F1 scores than other methods across the slides, followed by weighted Pearson and Spearman correlations, masked Spearman correlation, and finally bivariate Moran’s R (**Supp. Fig. S2A-B**).

Second, using 28 spatial transcriptomics slides from myogenic, ischemic, and fibrotic heart tissue upon myocardial infarction ^40^, we evaluated the ability of local ligand-receptor scores to recover cell type proportions (Methods). We noted that Cosine and Jaccard similarities had the highest predictive performance in ischemic (R^2^ ≈ 0.32) and fibrotic tissues (R^2^ ≈ 0.28), while Moran’s R did best in myogenic (R^2^ ≈ 0.13) (**Supp. Fig. S2C**); with similar results also observed in terms of Root Mean Squared Error (RMSE).

In summary, all spatially-informed local scores in LIANA+ preserved the biological signal of gene expression. Moreover, our results suggest that spatially-informed Cosine similarity, which performed best in both the regression and classification tasks, might be most suitable as a default local scoring function. However, the other scoring metrics are likely suited better for other tasks. For example, spatially-weighted Jaccard index should be well suited for categorical or binary data. Similarly, Spearman correlation should be more relevant when inferring relationships between ordinal, ranked, or non-linear variables. Thus, the choice of metric should take in consideration the data and task at hand.

### Supplementary Note 2. Sample Label Classification

To evaluate the ability of different combinations of ligand-receptor methods and dimensionality reductions to distinguish samples coming from different conditions, we set up a classification task (Methods). In each dataset, we inferred interactions independently for each sample using scoring functions from each ligand-receptor method in LIANA+, focusing on magnitude-based scores whenever available (**Supp. Table 2**). Then, we decomposed the ligand-receptor output, obtained per sample and method using MOFA+ and Tensor-cell2cell (**Supp. Fig. S5A**). Using a binary classification setup, we then calculate AUROC and F1 for each method-factorisation combination to see well each classified condition (**Supp. Fig. S5A**; Methods).

We saw that all combinations performed better than random in most datasets (**Supp. Fig. S5B**). On average all methods performed similarly when combined with MOFA+ (with average AUROCs between 0.83 and 0.85), with the exception of CellChat’s score (0.62 average AUROC; **Supp. Fig. S5B**).

On the other hand, we saw more variability between the methods when combined with Tensor-cell2cell. Specifically, ligand-receptor gene expression products, used by NATMI and Connectome, and LIANA’s consensus, with respective average AUROCs of 0.81 and 0.78, were most consistent across the datasets. The AUROCs of the rest of the methods ranged between 0.61 and 0.74 (**Supp. Fig. S5B**).

Moreover, MOFA+ had on average higher AUROCs than Tensor-cell2cell across all scores, except CellChat. This potentially reflects an intrinsic difference of the regularizations used between the two approaches. Specifically as MOFA+ attempts to enforce orthogonality ^51^, while the non-negative tensor component analysis used by Tensor-cell2cell, which can be thought of as a higher-order extension of NMF ^32^, does not. We also saw similar results when using weighted F1 (**Supp. Fig. S5C)**

Overall, our results show that both Tensor-cell2cell and MOFA+ are capable of capturing CCC events coordinated across cell-types that separate samples according to the different conditions at hand.

### Supplementary Note 3. LIANA+ enables joined CCC single-cell and spatial analysis

To jointly study CCC events in dissociated and spatially-resolved data, we used LIANA+ with two public datasets of mouse acute kidney injury (AKI) model ^52,53^. Both datasets employed a time course experimental design, in which murine kidneys were sequenced following bilateral ischemia-reperfusion injury (**Supp. Fig. S6A**).

First, using a single-nucleus AKI atlas (n=24) ^52^, we inferred potential ligand-receptor interactions between groups of cells at the sample level and decomposed the interactions with MOFA+ into a set of factors (Methods), with the aim to identify deregulated intercellular programmes associated with kidney injury. After quality control (Methods), we analysed CCC interactions across 88 cell type pairs and saw that Factor 1 separated early acute time points from the rest (Kruskal-Wallis P-value = 0.0069; **Supp. Fig. S6B**). Sample factor scores associated in Factor 1 were highest at the 12 hour time point, earlier than previously CCC results in a similar mouse model by Li et al., 2022 ^108^. Factor 1 explained on average 13.5% of the variability (R^2^) of ligand-receptor interactions across cell-type pairs, with Fibroblasts being involved in the several cell type pairs with variances explained > 30%, including their interaction as potential sources of communication with Proximal tubule epithelia (R^2^ = 52.7%; **Supp. Fig. S6C**). Fibroblasts were also the recipient cell type with the highest mean variance explained (average R^2^ = 23.5%; **Supp. Fig. S6C**), likely associated with their potential role in mediating the repair process following kidney injury ^109^.

Within the top 15 interactions associated with Factor 1 (**Supp. Fig. S6D),** we noted several potential interactions that involved Spp1 and Tnc, known to contribute to extracellular-matrix remodelling and tissue repair ^110,111^. Other extracellular-matrix interactions, such as Slit2 and Robo1/2, as well as Lama2 & Dag1, were negatively associated with Factor 1 (**Supp. Fig. S6D)**. Thus, Factor 1 potentially represents an intercellular response related to the disruption of the extracellular matrix and its remodelling.

To see if the interactions found between groups of dissociated cells are also captured in spatially-resolved data, we inferred potential interactions using LIANA+’s spatial component in five 10x Visium slides from the same AKI model^53^. We saw that the interactions between Spp1 and the Itgav/Itgb1 integrin complex increased both in spatial coverage, as well as co-clustering (Moran’s R) albeit low, subsequent AKI (**Supp. Fig. S6E**); in line with findings from Li et al., 2022 in dissociated single-cell data ^108^. Specifically, we saw that in the control slide, the interaction was localised mostly in a specific part of the kidney, the medulla, while subsequent to kidney injury the interaction was ubiquitous across the whole kidney, coherent with its ubiquitous Factor 1 loadings (**Supp. Fig. S6D**).

In summary, using LIANA+ we identified in a hypothesis-free manner intercellular programmes potentially involved in early response to AKI in dissociated single-cell data, and supported those using independent spatial transcriptomics samples.

---

**Supplementary Figure S1.**
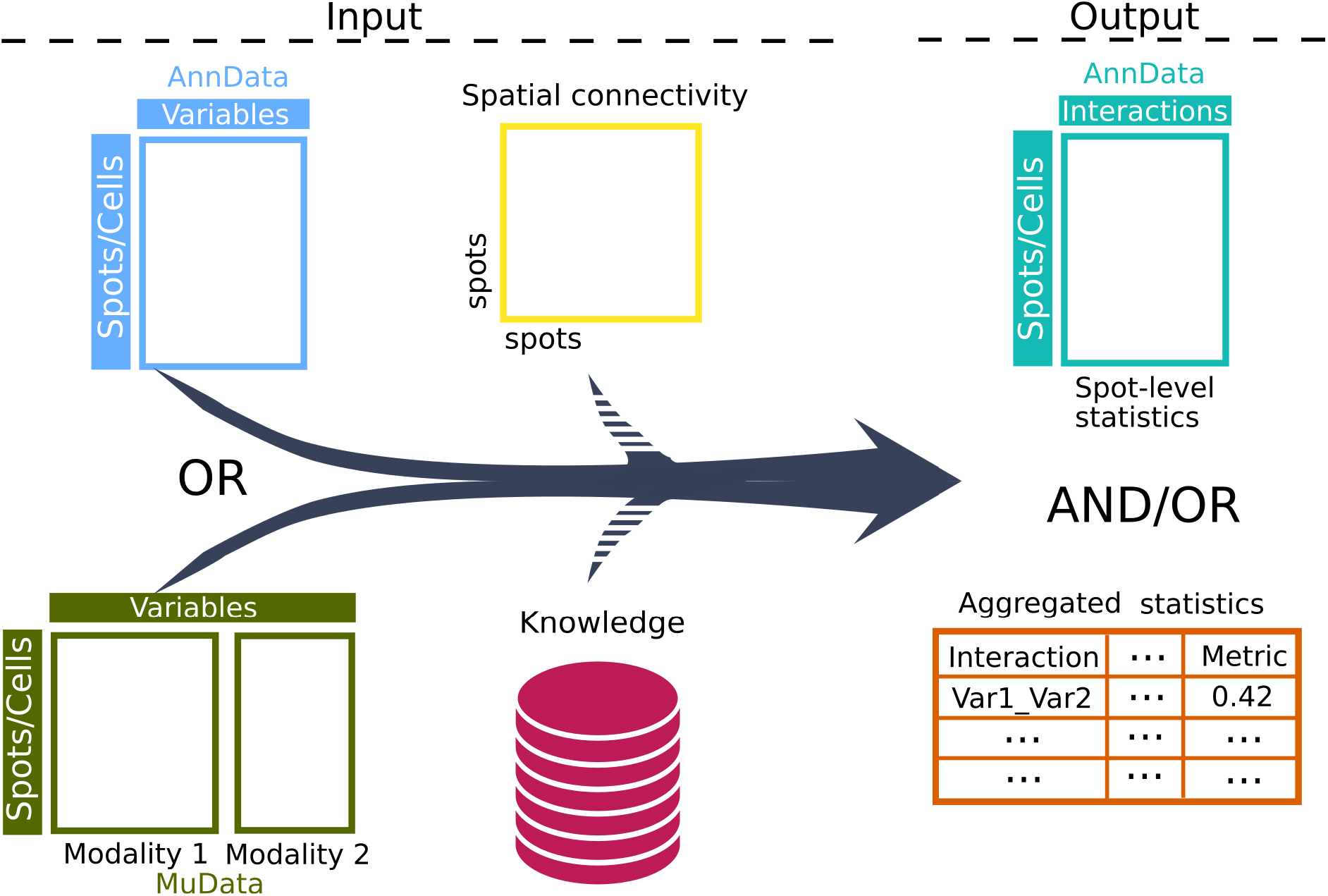
LIANA+ accepts inputs as unimodal (AnnData) or multimodal (MuData) data objects together with optional prior knowledge resource and/or spatial information. These are then transformed into dataframes of aggregated interaction results or statistics at the individual spot- or cell-level.

**Supplementary Figure S2.**
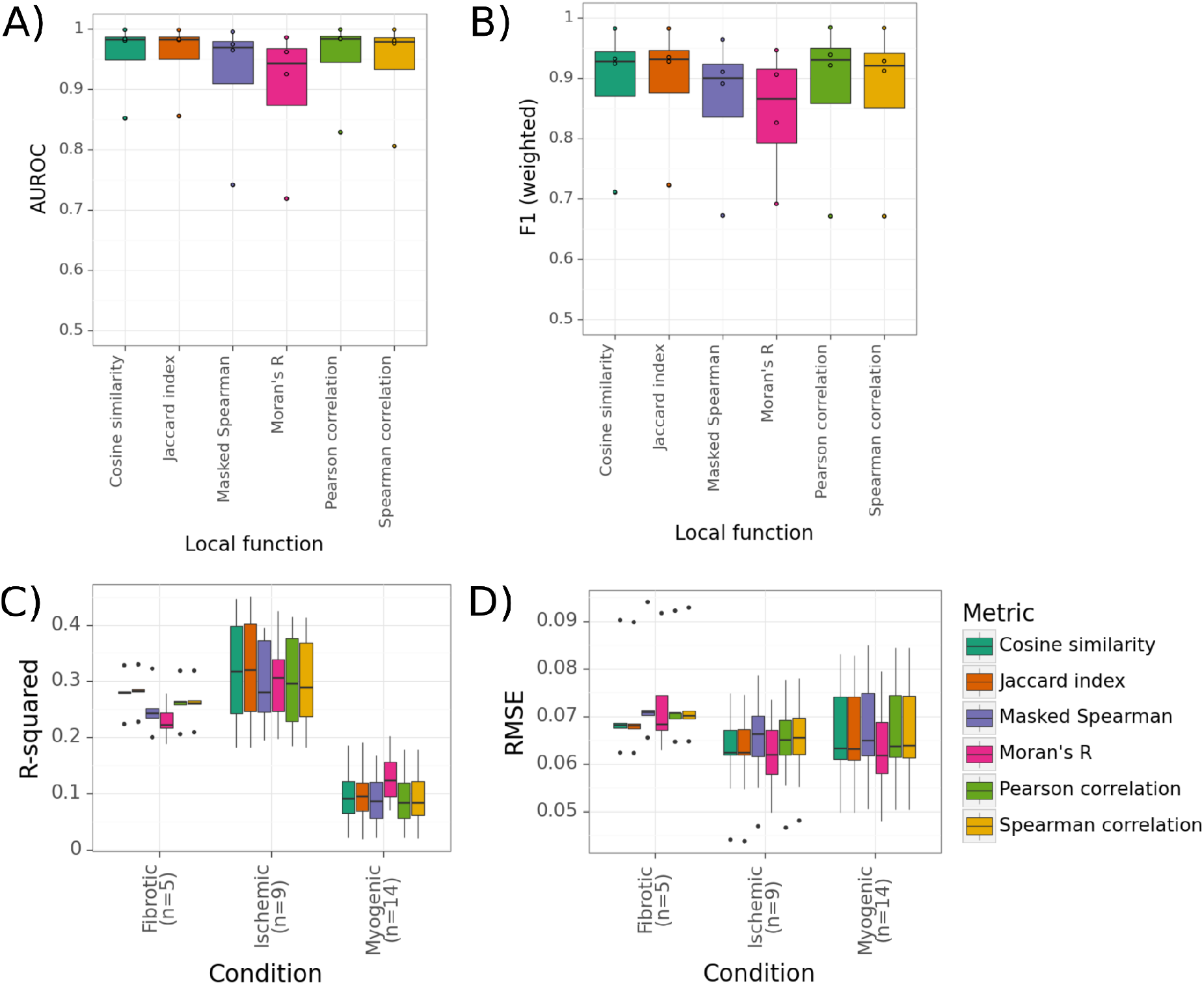
A) AUROC and B) weighted F1 for when using local metrics to classify malignant spots in breast cancer; C) R-squared and D) RMSE when using local metrics to predict cell type proportions in the heart.

**Supplementary Figure S3.**
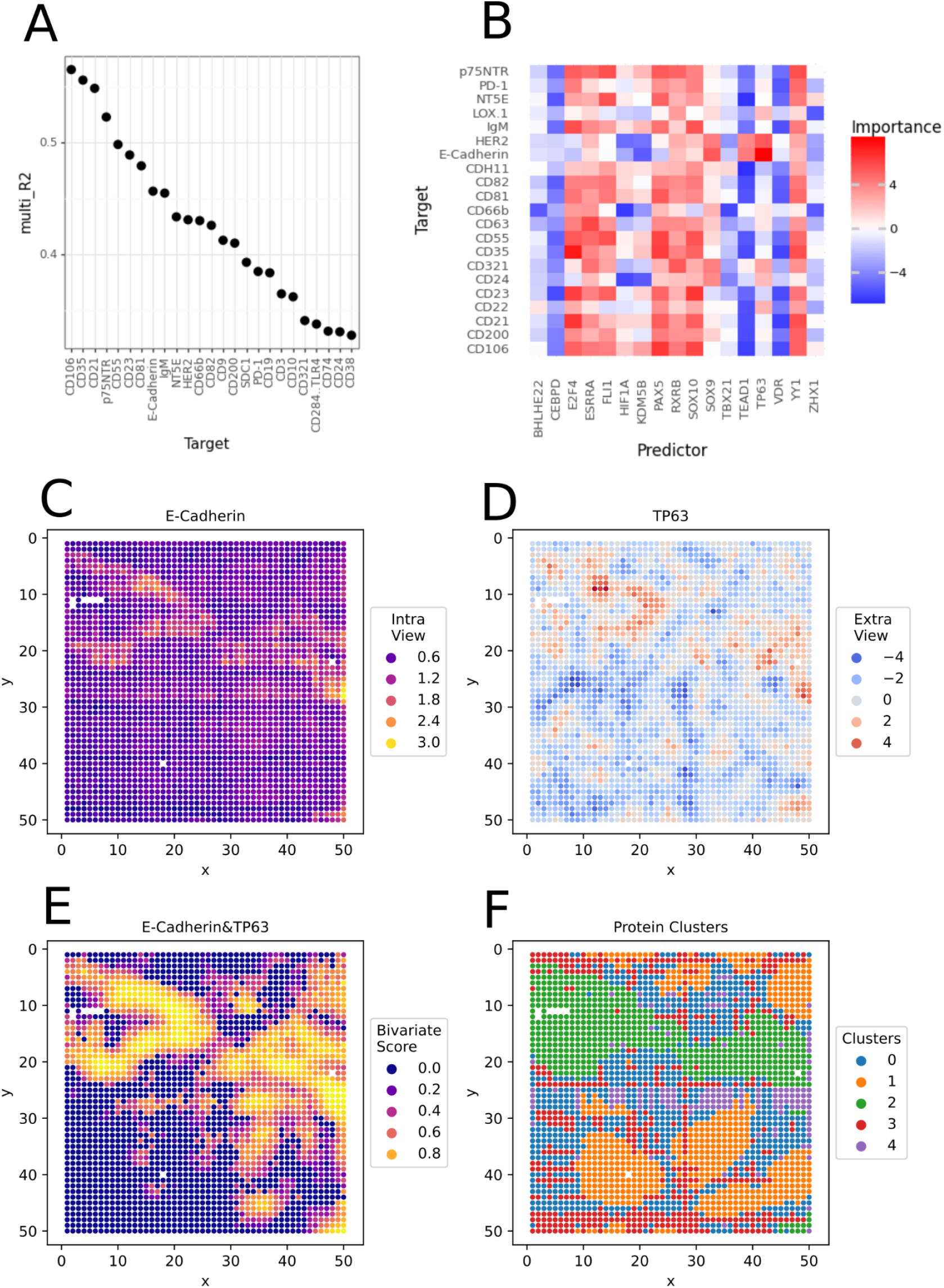
Analysis of spatial CITE-seq data from human secondary lymphoid (tonsil) tissue. A) Top 25 proteins with highest variance explained (R^2^), with 46% of the spatial variability of E-Cadherin being explained by transcription factor activities in neighbouring spots. B) Interaction scores (OLS t-values) for the top 50 interactions. C) E-Cadherin protein abundances. D) Spatially-smoothed TP63 transcription factor activity E) Spatially-weighted cosine similarity of E-Cadherin protein abundance and TP63 response. F) Spots clustered using the protein abundances.

**Supplementary Figure S4.**
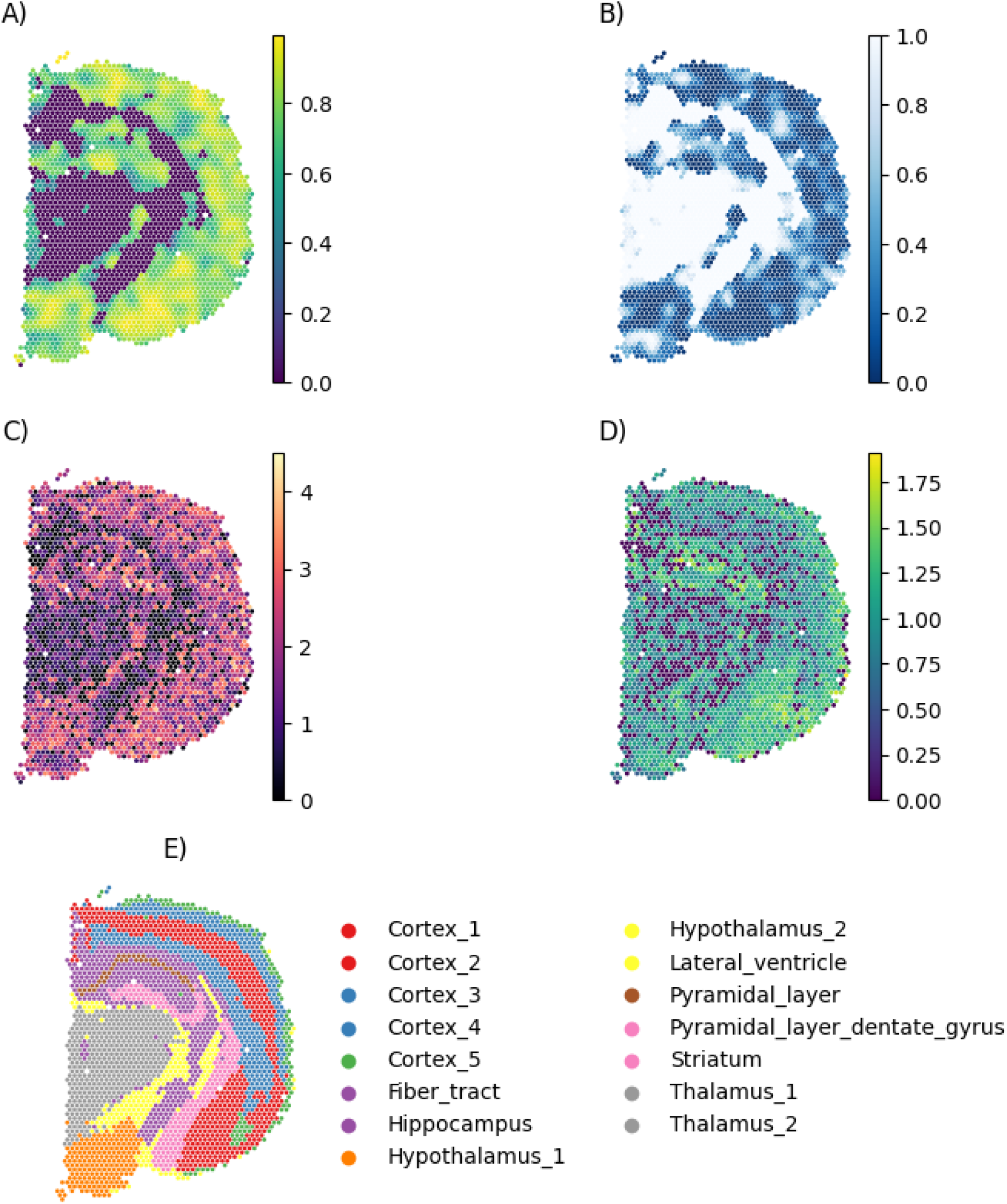
A) Local interaction between GABA & Gabra2, measured with Cosine similarity. B) Permutation-based local p-values. C) Estimated GABA abundances. D) Normalised Gabra2 Gene counts. F) Mouse brain niche annotations.

**Supplementary Figure S5.**
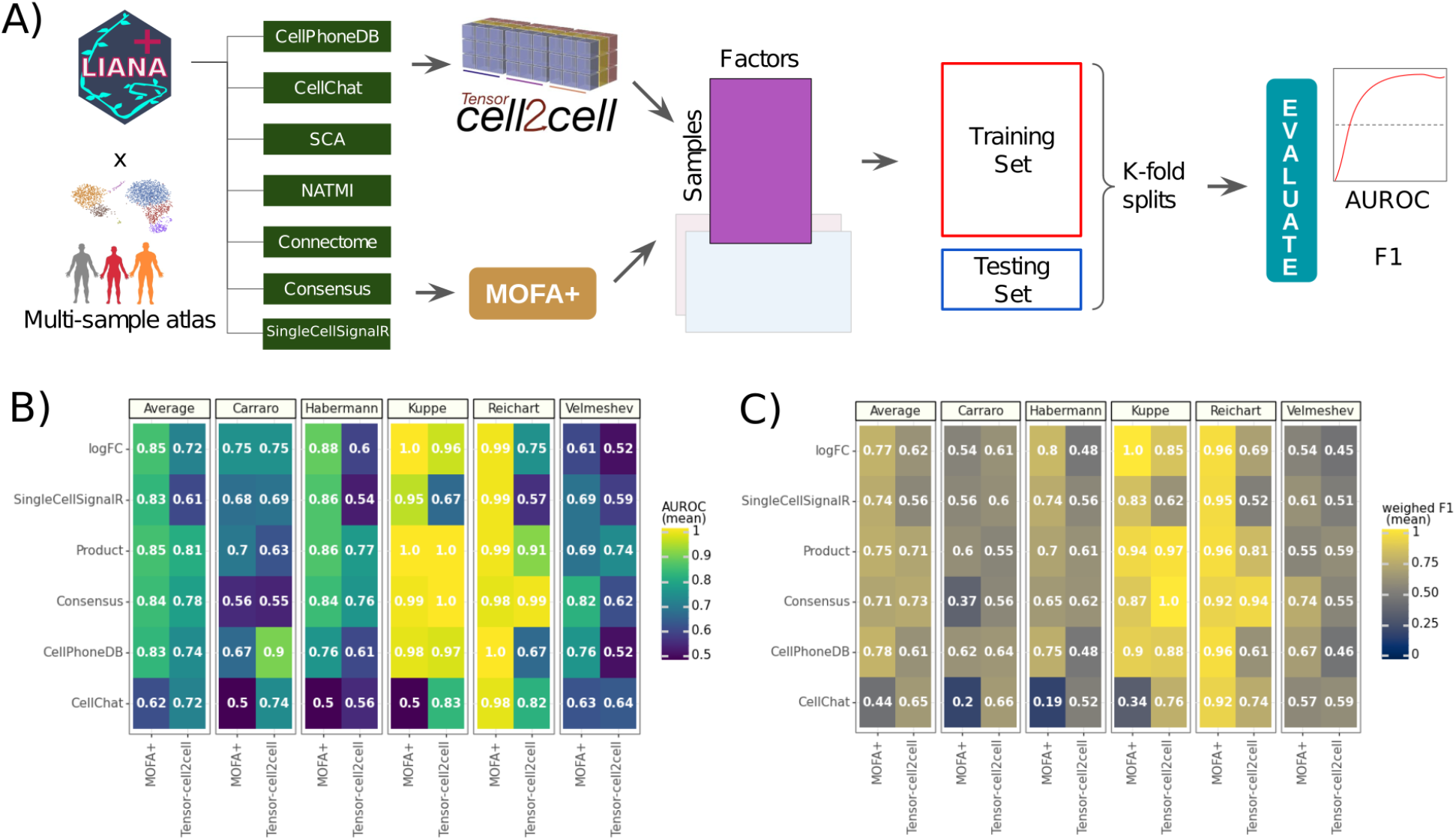
A) Classification setup to evaluate the ability of Tensor-cell2cell and MOFA+, combined with different ligand-receptor methods, to separate in an unsupervised manner conditions from multi-condition atlases. B) Average area under the receiver-operator curve (AUROC) and C) weighted F1 score.

**Supplementary Figure S6.**
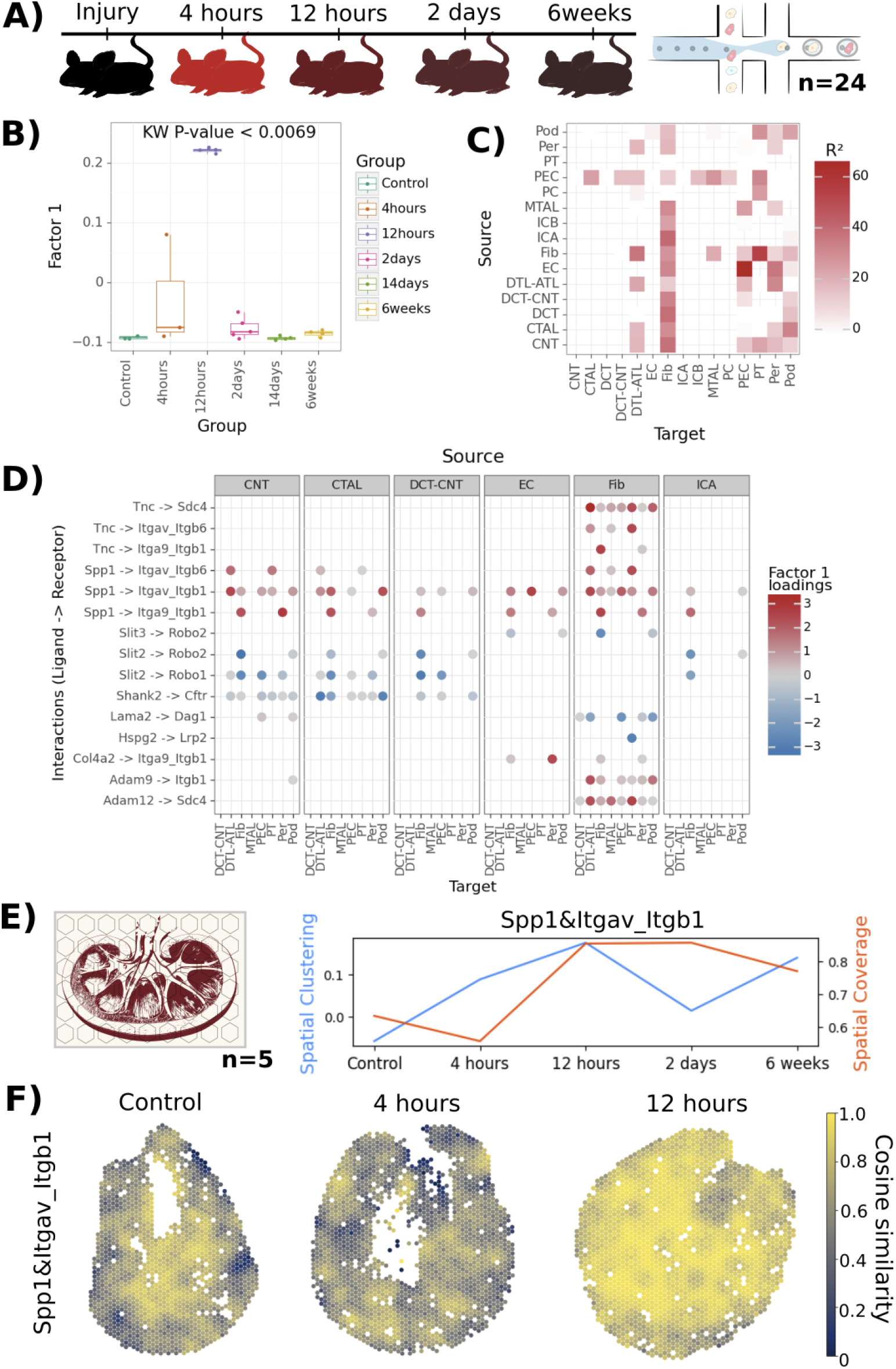
A) Experimental design of a murine AKI model ^52,53^. B) Distribution of Factor 1 sample scores at different time points following AKI. C) Variance explained by pairs of cell groups (views). D) Interaction loadings associated with Factor 1. E) Spatial Clustering (Global Moran’s R) and Coverage (mean Cosine similarity) of Spp1&Itgav_Itgb1 across conditions. F) Spatially-weighted Cosine similarity of Spp1 and the Itgav/Itgb1 complex in Control, 4 and 12 hours after injury.

**Supplementary Figure S7.**
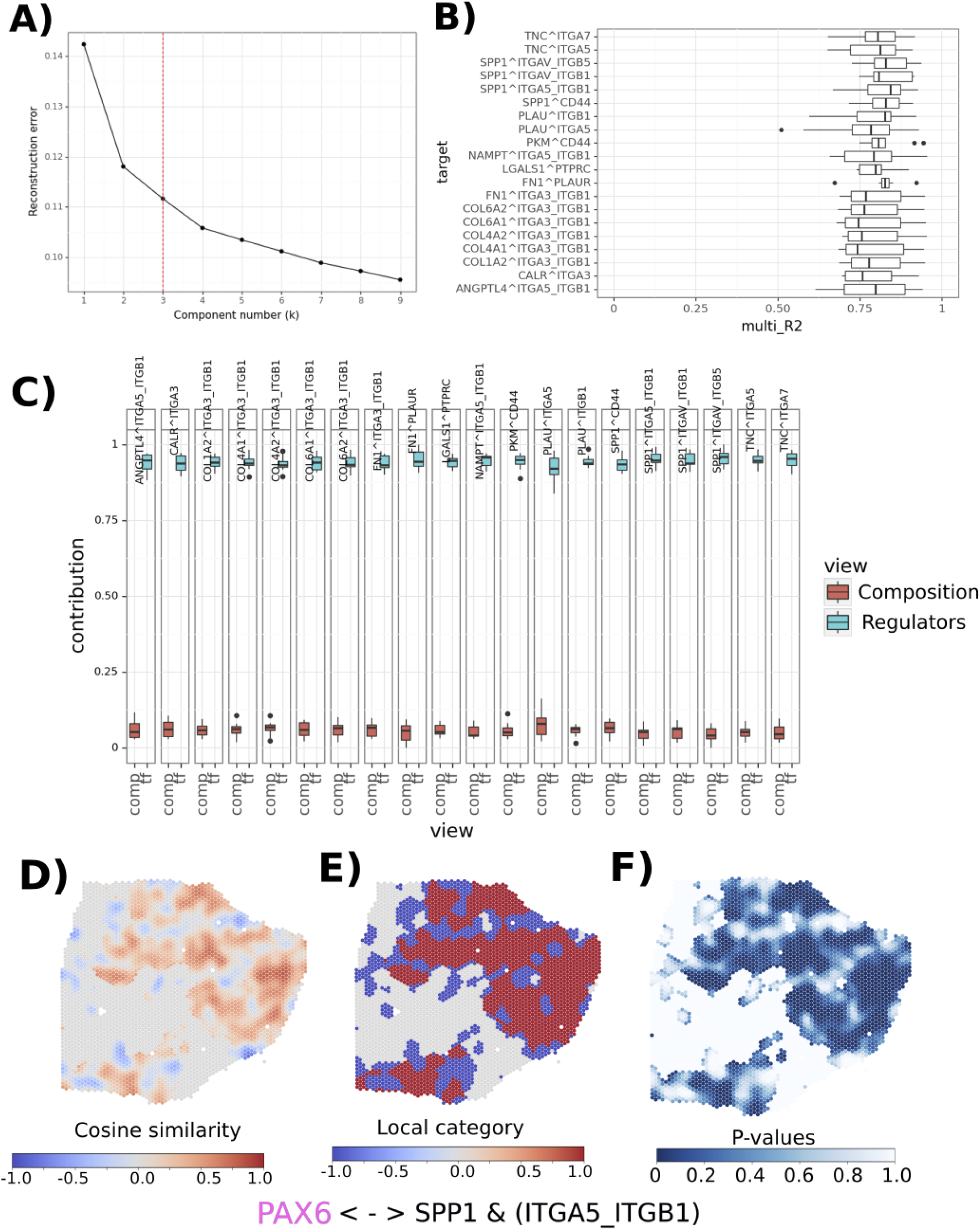
A) Elbow curve plot with a dashed red line showing the estimated optimal rank. B) Median R^2^ per Interaction. C) Contribution of Tissue Composition and Transcriptional Regulators. D) Cosine Similarity, E) Interaction category, and F) local permutation P-values for the spatial association between PAX6 and SPP1 & ITGA5_ITGB1.

**Supplementary table 1.**
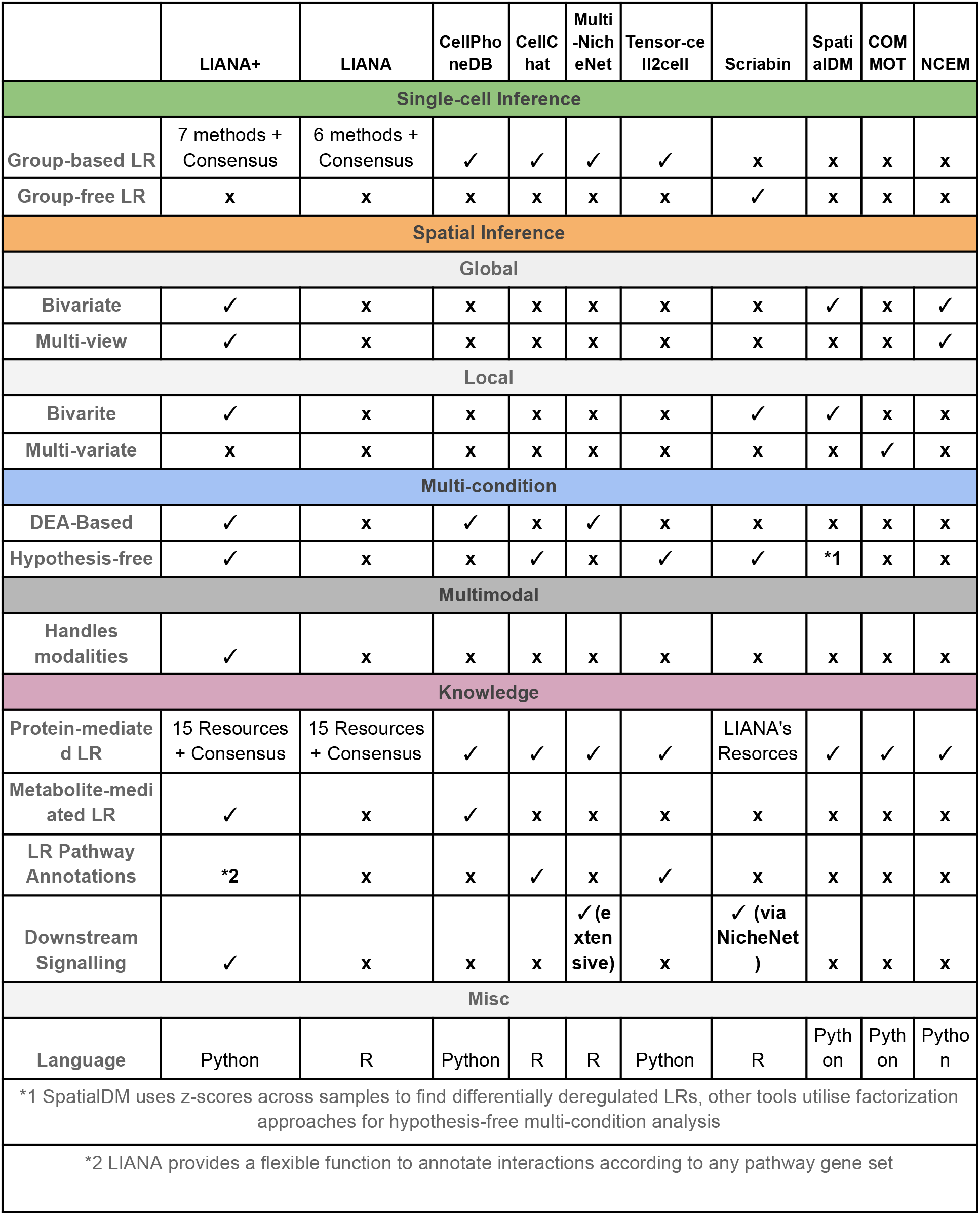
Feature comparison of selected CCC tools.

**Supplementary table 2.**
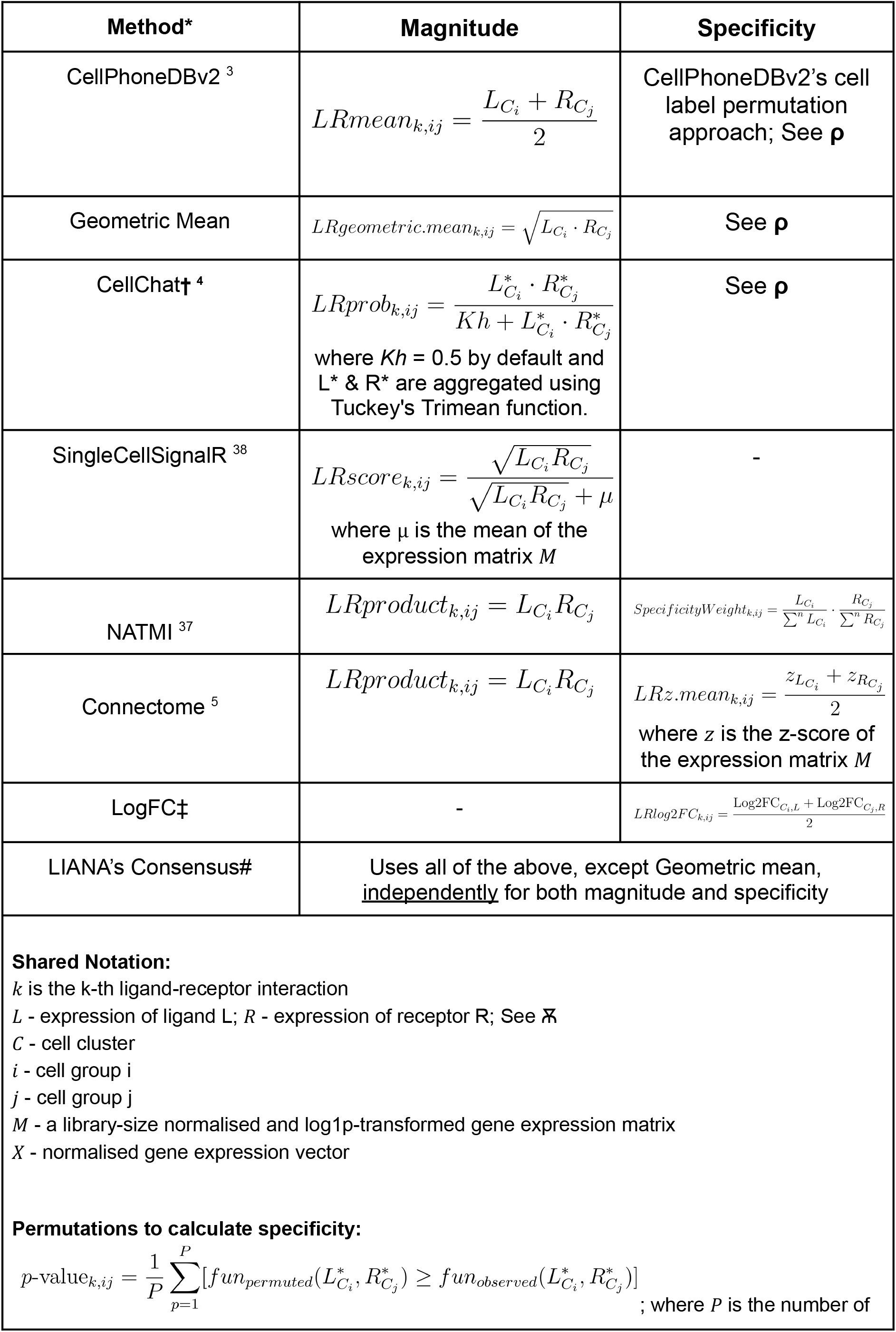
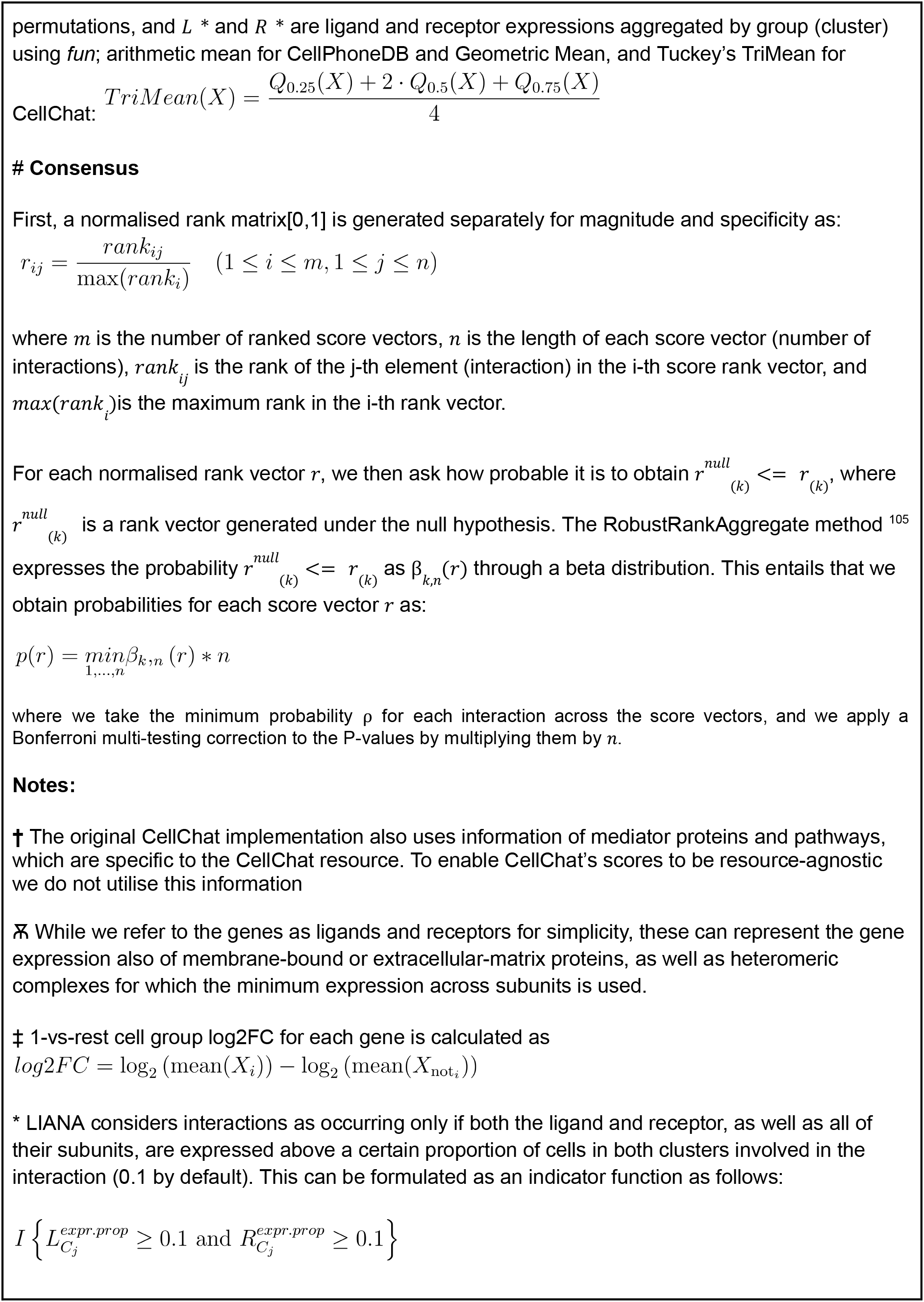
Single-cell ligand-receptor methods implemented in LIANA+.

**Supplementary table 3.**
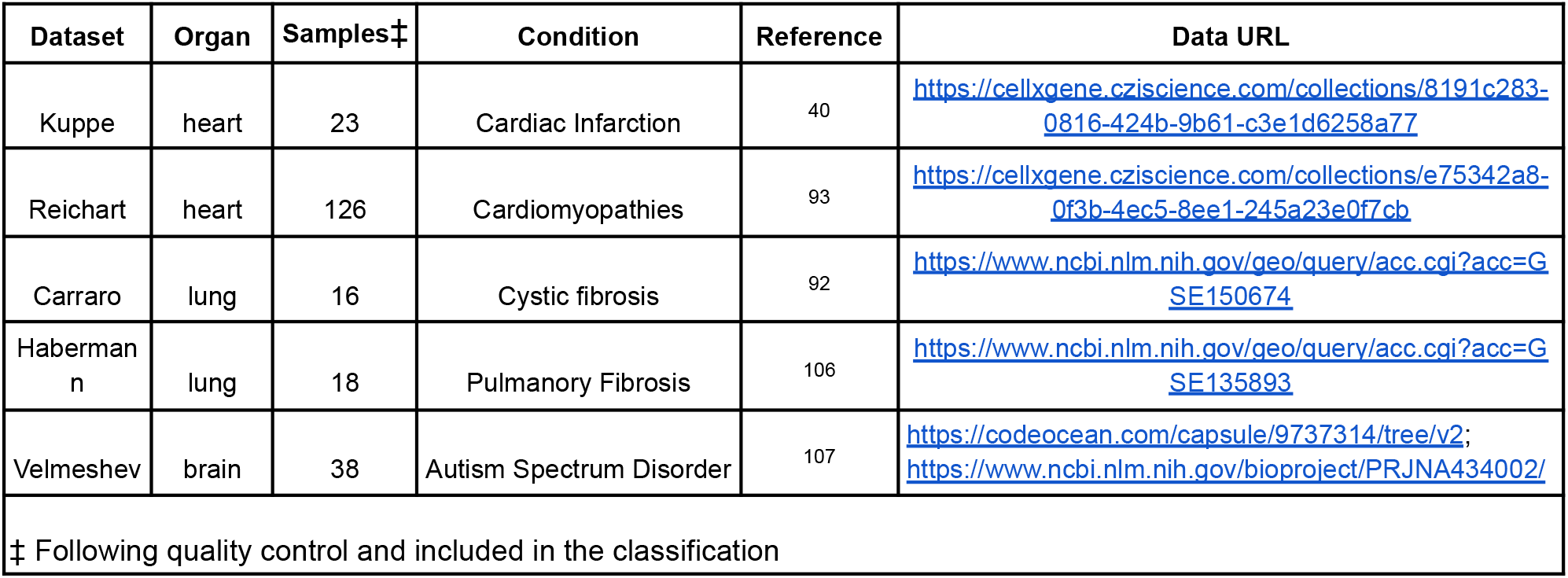
Cross-conditional atlases used in the sample classification task.

| Dataset | Organ | Samples‡ | Condition | Reference | Data URL |
| --- | --- | --- | --- | --- | --- |
| Kuppe | heart | 23 | Cardiac Infarction | 40 | https://cellxgene.cziscience.com/collections/8191c283-0816-424b-9b61-c3e1d6258a77 |
| Reichart | heart | 126 | Cardiomyopathies | 93 | https://cellxgene.cziscience.com/collections/e75342a8-0f3b-4ec5-8ee1-245a23e0f7cb |
| Carraro | lung | 16 | Cystic fibrosis | 92 | https://www.ncbi.nlm.nih.gov/geo/query/acc.cgi?acc=GSE150674 |
| Haberman<br>n | lung | 18 | Pulmonary Fibrosis | 106 | https://www.ncbi.nlm.nih.gov/geo/query/acc.cgi?acc=GSE135893 |
| Velmeshev | brain | 38 | Autism Spectrum Disorder | 107 | https://codeocean.com/capsule/9737314/tree/v2;<br>https://www.ncbi.nlm.nih.gov/bioproject/PRJNA434002/ |
| ‡ Following quality control and included in the classification |  |  |  |  |  |

